# Impact of sulfamethoxazole on a riverine microbiome

**DOI:** 10.1101/2021.04.01.438070

**Authors:** C. Borsetto, S. Raguideau, E. Travis, D.W. Kim, D.H. Lee, A. Bottrill, R. Stark, L. Song, J.C. Cha, J. Pearson, C. Quince, A.C. Singer, E.M.H. Wellington

## Abstract

The continued emergence of bacterial pathogens presenting antimicrobial resistance is widely recognised as a global health threat and recent attention focused on potential environmental reservoirs of antibiotic resistance genes (ARGs). Freshwater environments such as rivers represent a potential hotspot for ARGs and antibiotic resistant bacteria as they are receiving systems for effluent discharges from wastewater treatment plants (WWTPs). Effluent also contains low levels of different antimicrobials including antibiotics and biocides. Sulfonamides are antibacterial chemicals widely used in clinical, veterinary and agricultural settings and are frequently detected in sewage sludge and manure in addition to riverine ecosystems. The impact of such exposure on ARG prevalence and diversity is unknown, so the aim of this study was to investigate the release of a sub-lethal concentration of the sulfonamide compound sulfamethoxazole (SMX) on the river bacterial microbiome using a flume system. This system was a semi-natural *in vitro* flume using river water (30 L) and sediment with circulation to mimic river flow. A combination of ‘omics’ approaches were conducted to study the impact of SMX exposure on the microbiomes within the flumes. Metagenomic analysis showed that the addition of low concentrations of SMX (<4 μg L^−1^) had a limited effect on the bacterial resistome in the water fraction only, with no impact observed in the sediment. Metaproteomics did not show differences in ARGs expression with SMX exposure in water. Overall, the river bacterial community was resilient to short term exposure to sub-lethal concentrations of SMX which mimics the exposure such communities experience downstream of WWTPs throughout the year.

## Introduction

Antimicrobial resistance (AMR) has been extensively studied in clinical environments, but only recently non-clinical environments such as rivers have been recognised as potential hotspots for persistence and spread of antimicrobial resistance genes (ARGs) between bacterial species (Singer et al., 2016; Wellington et al., 2013). Rivers in the UK receive effluent discharges from wastewater treatment plants (WWTPs) and a significant prevalence of ARGs and antibiotic resistant bacteria (ARB) has been detected in rivers in close proximity to WWTPs (Brechet et al., 2014; Khan et al., 2019; Parnanen et al., 2019). ARGs are known to frequently transfer between bacteria and to be associated with mobile genetic elements (MGEs), which are useful markers to monitor pollution. Indeed, MGEs such as genes belongings to the class 1 integrons (*intI1*) were identified as a potential proxy for anthropogenic pollution in the environment (Amos et al., 2015; Gillings et al., 2015). In addition to ARB and ARGs, antibiotics are also released in the environment through wastewater effluent and their impact on the microbial community in the environment and spread of ARGs is still unknown. Currently no global standard regulations or environmental limits are in place for the control of antibiotic pollution and no additional treatment of wastewater effluent is performed unless required by local regulations (Carraro et al., 2016). The majority of antibiotics are not well metabolised in humans and animals; therefore they are excreted in the urine and reach the environment via WWTP discharges as well as via application of manure to land (Andersson and Hughes, 2012; Baietto et al., 2014; Byrne-Bailey et al., 2009; Kummerer, 2009). Antibiotic concentrations were reported ranging from 10 – 10^6^ ng L^−1^ in freshwater environments around the world, with low- and middle-income countries (LMICs) showing the highest concentrations reported for several antibiotics (Chow et al., 2021; Fekadu et al., 2019; Li et al., 2018). Studies have shown that even though β-lactams are the most widely used group of antibiotics in clinics (Gbaguidi-Haore et al., 2013; WHO, 2018), they are less detected in the environment due to breakdown by β-lactamase enzymes that are widely excreted by the microbial community (Brechet et al., 2014). However, other antibiotic groups such as fluoroquinolone, macrolide and sulfonamides, which are more stable, were reported at higher concentrations in wastewater effluents and rivers (Kummerer, 2009; Zhou et al., 2013).

Sulfonamides are frequently detected in freshwater worldwide with high concentrations reported for LMICs, where their consumption for clinical purposes is still prevalent (aus der Beek et al., 2016; K’Oreje K et al., 2016; Segura et al., 2015). These drugs target both Gram-positive and Gram-negative bacteria, affecting the dihydropteroate synthase (DHPS) enzymes involved in the folic acid biosynthesis (acts as an analogue of PABA thus binding DHPS), an essential precursor for the synthesis of nucleic acids (Skold, 2000). In the clinical setting, they have been used more frequently in combination with trimethoprim for the treatment of various urinary tract, respiratory, skin-associated and gastro-intestinal infections (Masters et al., 2003). Sulfonamides were used on a large scale in animal husbandry not only as treatment but also as animal feed additive, raising concerns for their effect on AMR selection (Baran et al., 2011; Grave et al., 2014). Regulations have therefore banned their use in animal feed in the last few decades, but they are retained for the treatment of animal disease (PHE, 2014). Resistance has been widely observed in the microbial community either through mutations of the chromosomal *dhps* (*folP*) gene or by acquisition of plasmid-borne genes (*sul1, sul2, sul3* and *sul4*) encoding for alternative drug-resistant versions of the DHPS enzymes which have lower affinity for sulfonamides (Achari et al., 1997; Razavi et al., 2017; Skold, 2000). In particular, *sul2* and *sul3* were reported on small non-conjugative plasmids or large transmissible multi-resistant plasmids (Enne et al., 2001; Perreten and Boerlin, 2003; Radstrom et al., 1991), while *sul1* and *sul4* were found in clinically relevant class 1 integrons typically associated with human pathogens (Gillings et al., 2008; Razavi et al., 2017).

Recent *in vitro* studies have shown that the presence of sub-lethal concentrations of antibiotics can select for AMR, suggesting that this effect could also be happening in natural environments where subinhibitory concentrations occur, thus promoting resistance in the natural microbial community (Gullberg et al., 2011; Stanton et al., 2020). The predicted no-effect concentration (PNEC) for resistance selection for sulfamethoxazole (SMX) varies widely from 0.6 to 16 μg L^−1^ (Bengtsson-Palme and Larsson, 2016; Le Page et al., 2018; Mortimer et al., 2020). Our aim was therefore to investigate the effect of a sub-lethal concentration of SMX in the PNEC range for selection of resistance on the microbial community of a riverine environment by use of *in vitro* flumes to mimic the effect of an antibiotic release event in a river ecosystem. Flumes have served as useful model systems to study pollutants in rivers (Cook et al., 2020a; Cook et al., 2020b). The focus was on SMX addition at a sub-lethal concentration in the range of previously recorded SMX measurements in rivers (Hanamoto et al., 2018; Vila-Costa et al., 2017). A combination of high-throughput qPCR and various ‘omics’ were used to demonstrate that the riverine microbiome was resilient to sublethal challenges with SMX <4 μg L^−1^.

## Methods

### Flumes set up

River water and sediment were collected downstream of a WWTP form the River Sowe, Stoneleigh, UK in April 2019 and immediately used to set up the flumes which consisted of six individual flumes, each containing 6 kg of sediment (3 cm depth river bed) and 30 L of water (12 cm water column over the river bed) to replicate the river environment. As previously described by (Cook et al., 2020b), each flume (2.36 m length × 0.2 m height × 0.1 m width) was made of glass and was connected to a Haillea HC-300A aquarium chiller (Hailea Group Co., China) to keep the temperature system constant to 20 °C and a pump to mimic the river flow (10 L min^−1^) monitored by a GPI TM Series electronic flow meter (Great Plains Industries, Inc., US). Flumes were covered with clear polythene sheets to limit evaporation. A light cycle was set to 16 h light and 8 h dark using a fluorescent 70 W daylight bulbs (F70W/865 T8 6ft, Fusion Lamps, UK) with LEE226 filters (Transformation Tubes, UK).

After set-up, the system was left to settle for 36 h before starting the flow and allowed to equilibrate for an additional 24 h. After this initial settling phase, 3 flumes were amended with SMX at 4 μg L^−1^, while the other 3 flumes were left untreated (controls). The experiment was run for a total of 24 days and samples (water and sediment) were taken from each individual flume at selected time points (0, 3, 7, 12, 18, 24 days). For time point 0, samples from the amended flumes were collected after 1 h of addition of the SMX in order to allow the compound to evenly distribute in the system.

### Chemical analyses

Chemical analyses were performed only on the water samples throughout the experiment to monitor organic C, NH_4_ and inorganic ions NO_3_, NO_2_, pH, alkalinity, dissolved oxygen and SMX concentration. For the latter, 2 mL of water were filtered (0.22 μm PES filter) and liquid chromatography analysis performed with Waters H-class UPLC and Waters Xevo TQXS triple quad mass spectrometer using a 50 × 2.1 mm C18 Waters X-bridge column with particle size of 1.7 μm. The mobile phases consisted of water with 0.1% formic acid (A) and acetonitrile with 0.1 % formic acid (B). The run gradient was performed with a flow rate of 0.4 mL min^−1^ following the conditions: 0-1 min 80 % A – 20 % B; 1-3 min 5 % A – 95 % B; 3-4 min 95 % B; 4-5 min 80 % A – 20 % B. The column was equilibrated for 3 min before the next injection. The MRM transitions was set to 254.12>91.98 (CE 32eV), 254.12>98.94 (CE 16eV). The SMX isotope (phenyl-13C6, 99%) (Alpha isotopes limited) was used as internal standard to quantify the SMX concentration.

### Bacterial community analysis

Water (0.1-1.5 L) and sediment (20-30 g) samples were collected from each individual flume. Water samples were immediately filtered (0.22 μm PES filter) and the filters were stored in sterile petri dishes at −20 °C until processed. Sediment samples were immediately stored at −20 °C until extraction. FastDNA™ Spin Kit (MP Biomedicals™) was used to extract DNA from 0.5 g of sediment and 0.5 L of water (0.22 μm PES filter). All DNA samples were quantified by Qubit (ThermoFisher) and quality check was performed by Nanodrop spectrophotometer (ThermoFisher) and gel electrophoresis prior to any further application. DNA was used for 16S rRNA V3-V4 amplicon sequencing using 300 bp paired-end Illumina Miseq. For selected time points (0, 7 and 18 days) the Wafergen SmartChip high-throughput qPCR with Primer set 2.0 was performed to quantify a total of 380 ARG and MGE targets on each sample (Stedtfeld et al., 2018). Metagenomes (average 90M-130M raw reads = 27 Gbp-37 Gbp × water and sediment samples respectively) were sequenced for selected timepoints (0, 7 and 18 days). Libraries were prepared and sequenced by Novogene (PE 150 bp – Illumina HiSeq).

Metaproteomic extraction was performed on selected water samples (time points 0, 7 and 18 days) using an optimized version of the protocol described by (Colatriano and Walsh, 2015). Briefly, cells from 0.5 L of water were collected on a 0.22 μm PES filter which was incubated in SDS lysis buffer (Tris-HCl 0.1 M pH 7.5, glycerol 5%, EDTA 10mM pH 8, SDS 1 %) for 20 min with agitation. Samples were then boiled for 20 min and incubated for 1 h at room temperature before centrifugation at 4000 rpm 10 min. The supernatant was transferred to 10 KDa Amicon Ultra tubes (Merck) and the boiled filter was washed with fresh SDS lysis buffer, then added to the protein sample. The supernatants were concentrated using 10 KDa Amicon Ultra membrane and the retained samples were washed twice with fresh SDS lysis buffer. A solution of methanol:acetone (50:50) was added to the concentrated samples in a 4:1 (vol/vol) proportion and protein precipitation was performed ON at −20 °C. The precipitated proteins were recovered by centrifugation at 17000 g for 30 min, dried by speedvac and resuspended in the SDS lysis buffer. Proteins were quantified using the BCA Pierce kit (ThermoFisher) and equal amount of proteins per sample (13 μg) were loaded on a pre-casted RunBlue™ TEO-Tricine gel (Expedeon) in presence of LDS and DTT. The gel was run for 10 min 180 V and stained with InstantBlue™ (Expedeon). Gel bands were cut and in-gel digestion with Trypsin was performed following reduction and alkylation with TCEP and CAA. Peptides were eluted from the gel and resuspended in 2 % acetonitrile + 0.1 % formic acid. Peptide extracts were analysed by nanoLC-ESI-MS/MS using the Ultimate 3000/Orbitrap Fusion instrumentation (Thermo Scientific) with a 60 min LC separation on a 25 cm column.

### Data filtering and statistical analysis

#### Chemicals

Statistical analysis was performed in R (v. 3.6.0). Package ggplot2 was used for graphical visualization.

#### Smartchip high-throughput qPCR

For each sample three technical replicate were run and for each condition three biological samples were analyzed. The resulting qPCR dataset was filtered according to Ct values threshold of 31, showing an efficiency between 1.7 and 2.3 and removing data showing Ct outlier and multi melt peaks as previously reported (Chen et al., 2019; Wang et al., 2014). Additionally, only technical replicates with 2 and 3 positive amplification were kept for further analysis. For biological replicates, only targets that had positive amplification for all 3 replicates were used for analysis. ARG data were normalized against 16S rRNA gene copy number. Data filtration was performed in Microsoft Excel with statistical analysis and graphical representation using R (v. 3.6.0).

#### 16S rRNA gene amplicon

Data were processed using the Qiime2 platform with DADA2 for denoising and ASV calling and Silva database for taxonomic annotation. Samples were rarefied at 23700 sequences per samples. Qiime2 and R (v. 3.6.0) packages phyloseq, vegan and NMIT were used for diversity and statistical analysis and ggplot for plots generation.

#### Metagenomes

general bioinformatic analysis were provided by Novogene according to the company’s pipeline as follows: all metagenomes raw data were quality filtered, then assembly of single sample metagenome and mixed assembly of the unutilized reads of each sample were performed using MEGAHIT (k-mer = 55). The gene prediction was carried out with MetaGeneMark (v 2.10) based on the scaftigs which were assembled by single and mixed samples. Pool predicted genes were dereplicated (CD-HIT v.4.5.8, identity = 95 %, coverage = 95 %) to generate gene catalogues and gene abundance in each sample was calculated by total number of mapped reads to gene catalogues (SoapAligner v 2.21). Reads taxonomy annotation was performed against the database of taxonomically informative gene families (NR database), while functional annotation was inferred based on similarity to sequences in the KEGG, eggNOG, CAZy and CARD (blastp, evalue ≤ 1e-5) databases.

Taxonomy and functional composition were explored using clustering analysis, Anosim, PCA, NMDS, Metastats and LEfSe multivariate statistical and comparative analysis based on abundance tables. Additionally, for the water metagenomes only, Metagenome Assembled Genomes (MAGs) were also recovered. MAGs taxonomic annotation was performed against the gdtb database for taxonomic reference.

#### Metaproteome

Water metaproteomes spectra were matched to the protein sequences present in the Comprehensive Antibiotic Resistance Database (CARD v 3.0.9) using MaxQuant (v 1.6.7.0). Perseus (v 1.6.2.2) was used for statistical analysis of the label-free quantitative (LFQ) values.

## Results

### Flumes chemical analysis

During the 24 days experiment, the nutrient and chemical properties within the water flumes were tested (pH, gran alkalinity, N, NH_4_, NO_2_, NO_3_, F, Cl, SO_4_, C, O_2_). No significant differences were observed in presence of SMX with the exceptions of dissolved sulphate (Kruskal-Wallis, p < 0.001) and chloride (Kruskal-Wallis, p < 0.01) which declined more rapidly in the presence of the antibiotic and the alkalinity which increased over time in presence of SMX (Kruskal-Wallis, p < 0.05) (Fig S1).

The initial concentration of SMX in the control river water was 127±30 ng L^−1^, while the amended flumes showed a concentration 10X higher with 1219±584 ng L^−1^ after the addition of the antibiotic. A rapid decrease in the level of SMX was observed after 3 days with a return to an equal or higher concentration than the initial one after just 7 days (Fig S1). This pattern in the first 7 days has been observed on several occasions in this system and in similar scaled down systems (data not reported). After the initial 7 days the SMX concentration decreased gradually.

### Microbiome structure

Alpha diversity analysis of 16S rRNA gene amplicon sequencing data showed a lower diversity for water samples compared to the sediment (Fig S2). The Chao1 richness index ranged from 2088 to 843 in water, whereas it was significantly higher in sediment where it ranged from 3951 to 1825 (Kruskal-Wallis H=51.75, p-value<8.5e^−13^). Similarly, the water samples also showed a significantly lower evenness with a Shannon index between 3.8 and 6.0 compared to the sediment samples between 6.9 and 7.4 (Kruskal-Wallis H=52.5, p-value<2.005e^−13^). The microbiome diversity in both environments did not significantly change over time throughout the experiment (Kruskal-Wallis, p-value>0.05) nor were they significantly affected by the presence of SMX (Kruskal-Wallis, p-value>0.05). Within the top 10 represented phyla, Proteobacteria was the most abundant phylum in both environments (50.2±8.7% in water and 33.4±1.6% in sediment) (Fig S3). In sediment the second most abundant phylum was Chloroflexi (15.8±1.1%), followed by Bacteroidetes (10.5±1.4), Actinobacteria (9.5±1.4%) and Firmicutes (6.2±1.0%). In the water column the second and third most abundant phyla were represented by Bacteroidetes (15.2±10.5%) and Planctomycetes (6.0±3.3%) followed by Actinobacteria (4.7±3.2%) and Nitrospirae (4.7±4.5%) (Fig S3). Beta diversity analysis based on Bray-Curtis dissimilarity at the ASV level showed a clear separation between the community composition present in the water and the sediment samples (Adonis, R2=0.36, p=0.001) (Fig. 1A).

**Fig 1.**
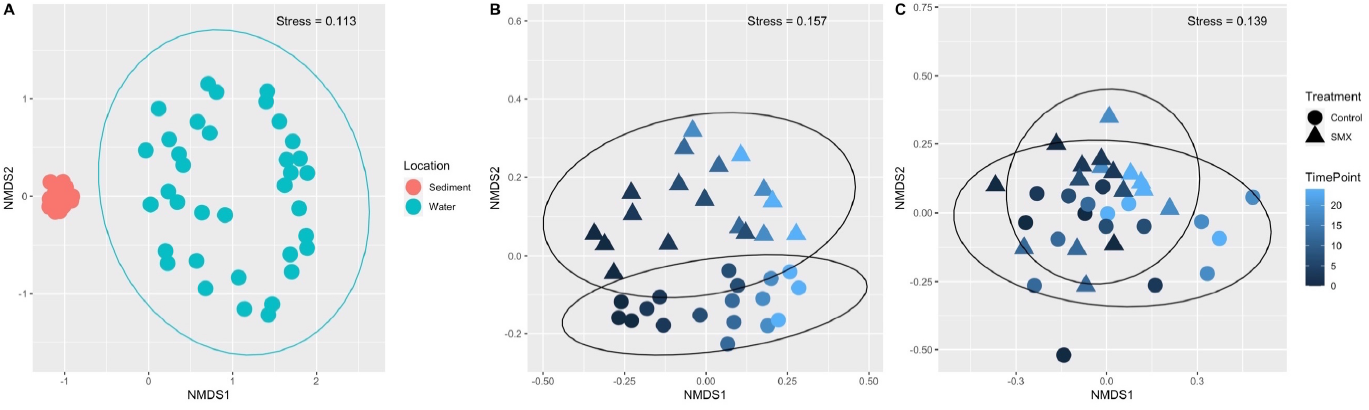
Microbial community ordination by NMDS of Bray-Curtis distance based on 16S rRNA amplicon sequencing data. A) All sediment and water samples – grouping by Location; B) Water samples - grouping by Treatment (control/SMX); C) sediment samples – grouping by Treatment (control/SMX). In B and C different time points are represented by colour changes, while Treatment by shape.

In particular, in the water the microbiome compositions varied across the flumes (Adonis, R2=0.28, p=0.001) and both time and treatment impacted equally on the community diversity, showing grouping according to treatment (Adonis, R2=0.14, p=0.001) and time (Adonis, R2=0.14, p=0.001) (Fig 1B). The sediment microbiome was more consistent throughout the experiment (Adonis, R2=0.06, p=0.002) and in the presence of SMX (Adonis, R2=0.05, p=0.004) (Fig 1C). Comparable results were also obtained when similarity and phylogenetic relationship between communities were tested using the Jaccard and weighted Unifrac indices (Fig S4), showing that the microbial community in the sediment was less affected by experimental variations and the presence of SMX compared to the water microbiome. However, to further evaluate the temporal interdependence between taxa amongst the control and SMX amended groups, nonparametric microbial interdependence analyses which used temporal correlation between taxa were also explored. These analyses at the same ASV level showed that 31 % of the microbial community variation over time was explained by the presence of SMX in the water (Adonis R2= 0.31), while only 22 % was attributed to SMX in the sediment (Adonis, R2= 0.22) (Fig S5). Although the microbial interdependence profiles between control and SMX groups were not statistically significant (p=0.1), the results suggest that sediment microbiome was more resilient to changes than the water microbiome. Similarly, at the metagenome level, stronger correlation coefficients were observed between sediment samples compared to the water samples (Fig S6).

Differential analysis of the water metagenomes at genus level identified potential general microbial biomarkers characterizing both control and SMX microbiomes throughout time. For instance, the abundance of genera such as *Nitrospira* and *Legionella* significantly increased after 7 days in both control and SMX groups overtime, while *Flavobacterium, Lysobacter, Phenylobacterium, Polynucleobacter* and *Methylopumilus* decreased (Fig 2).

**Fig 2.**
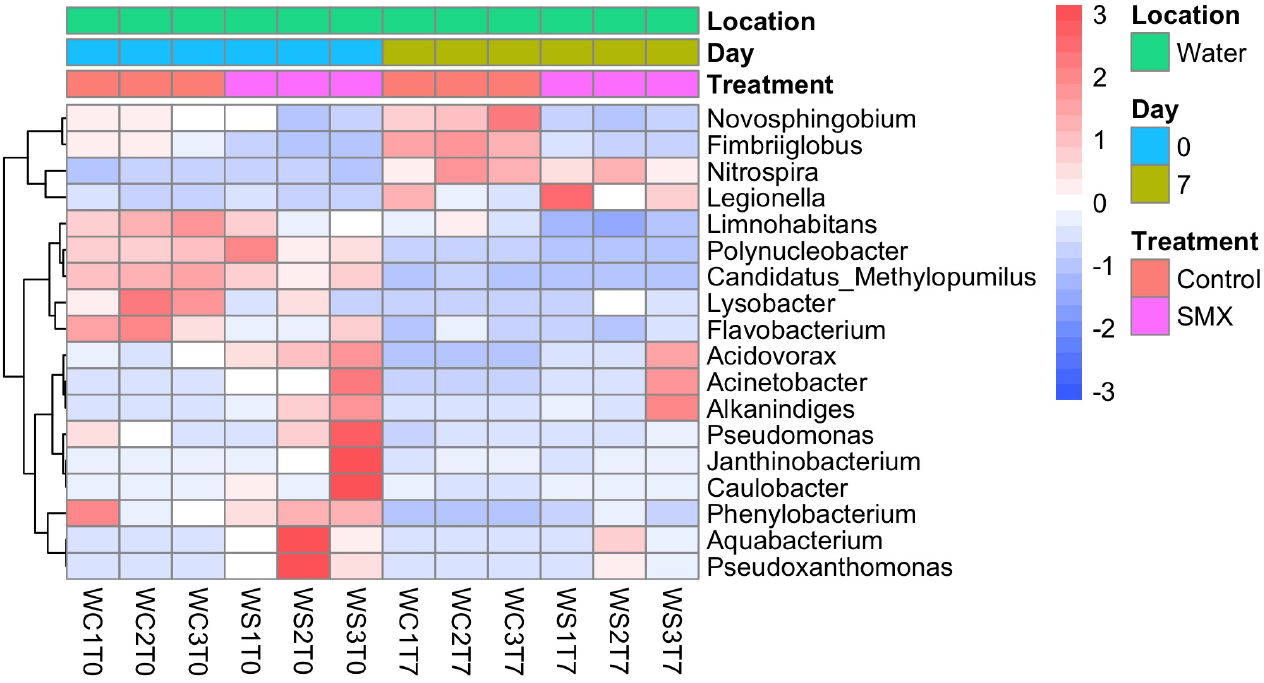
Distribution of biomarkers (Genus level) identified by LEfSe analysis in water samples metagenomes. Standardized Z values of relative abundance are represented for each selected biomarker genus for Control and SMX groups at time point 0 and 7 days.

### Microbiome functional analysis in response to SMX

The microbial functional profile investigated in the metagenome through annotation against the CAZy database indicated that the presence of SMX did not significantly affect the microbiome carbohydrate metabolic pathways. However, it was observed that in the water samples the carbohydrate utilization pathways changed over time in both control and SMX groups with an enrichment of the microbiome towards microorganisms containing genes related to the breakdown and modification of oligosaccharides rather than the metabolism of more complex polysaccharides (Fig S7).

The investigation of unique gene functionality by KEGG pathways assignation showed that 2.65 % of the unique genes of all samples were involved in the biodegradation and metabolism of xenobiotics and 1.61 % were assigned to pathways involved in drug resistance to antimicrobials (Fig S8). In particular, both pathways were more represented in the sediment samples (Fig 3).

**Fig 3.**
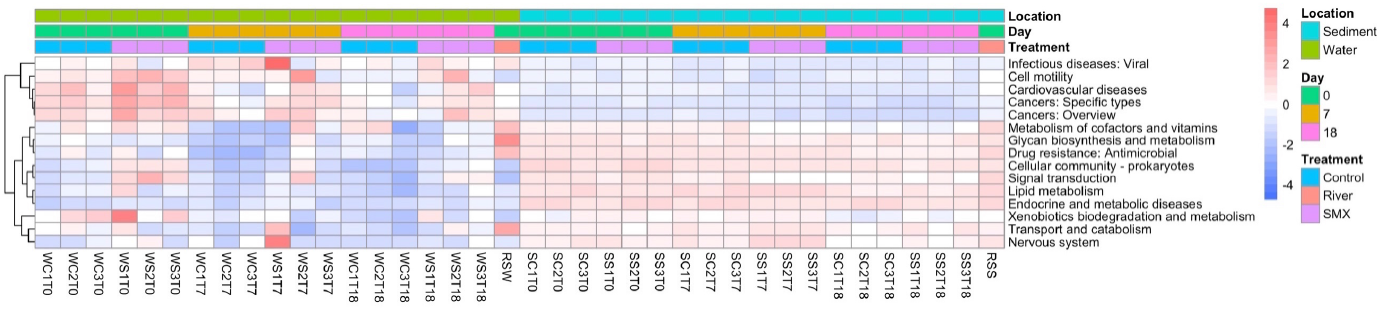
Functional KEGG domains characterising water and sediment microbiomes. Identification of characterising pathways by MetaStats analysis of metagenomic reads. Standardized Z values of KEGG domains abundance are represented.

A more detailed analysis of the metagenomes’ functional genes related to drug resistance to antimicrobials using the CARD database identified ARGs across both water and sediment, with efflux pumps conferring multi-drugs resistance being the most abundant class followed by fluoroquinolone and aminoglycoside (Fig. 4).

**Fig 4.**
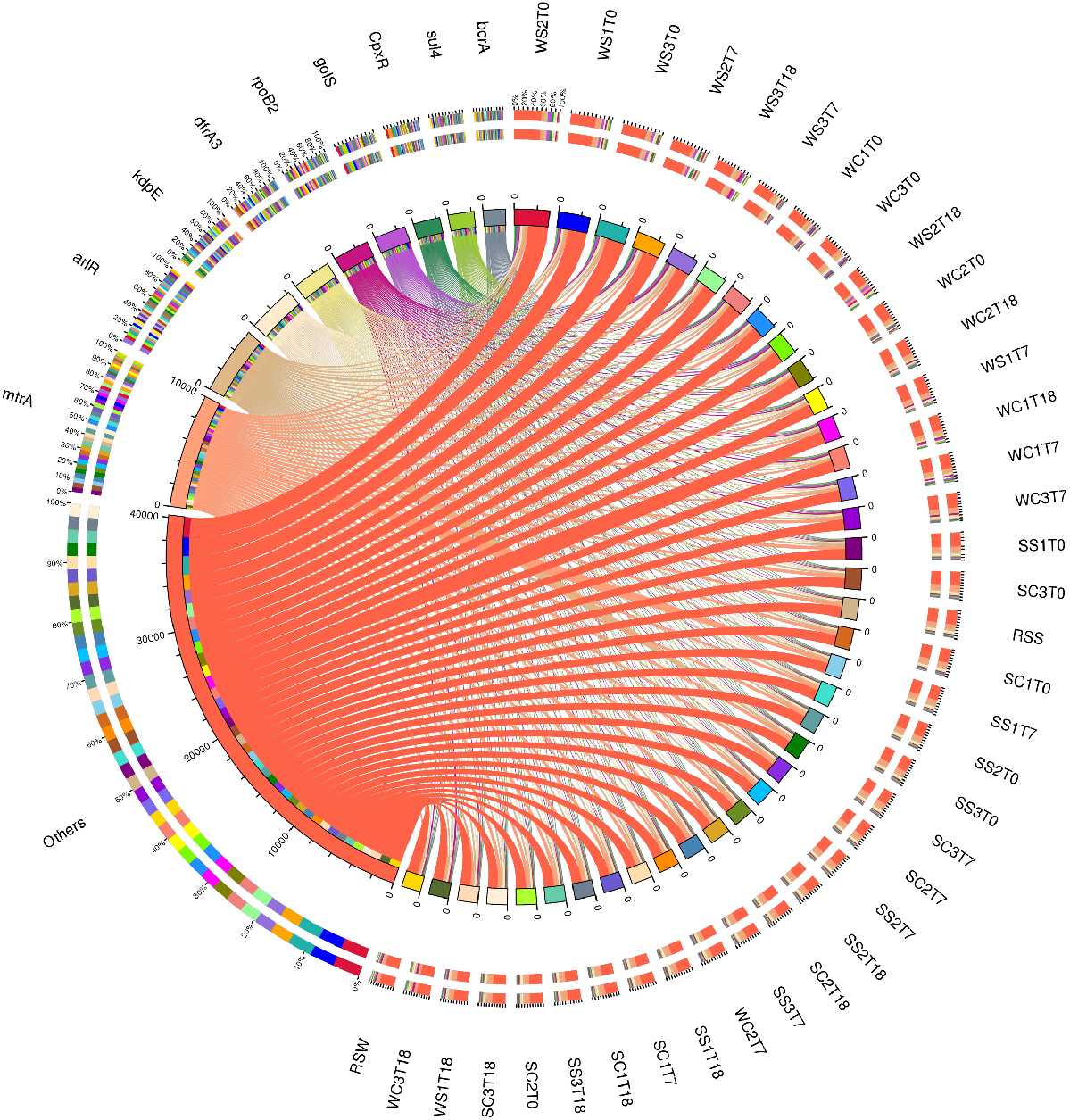
ARG distribution across water and sediment metagenomes. Samples are represented on the right side of the circle and ARGs on the left side. Inner circle: different colours represent different samples and ARG; the scale represents the relative abundance (ppm). The left part represents the sum of relative abundance of different samples in ARGs, while the right side is the sum of the relative abundance of the ARGs in each sample. Outer circle: the left part represents the relative percentage composition of different samples in ARGs, the right part vice versa.

Significant differences in the relative abundance of ARG classes were observed between water and sediment metagenomes (Adonis R2=0.41, p<0.05). For both microbiomes, significant changes were observed overtime throughout the experiment (Adonis: water R2=0.2, p=0.007; sediment R2=0.29, p=0.001), but the presence of SMX only had a significant impact on the ARG prevalence for the water microbiome (Adonis R2= 0.31, p=0.001) (Table 1).

**Table 1.**
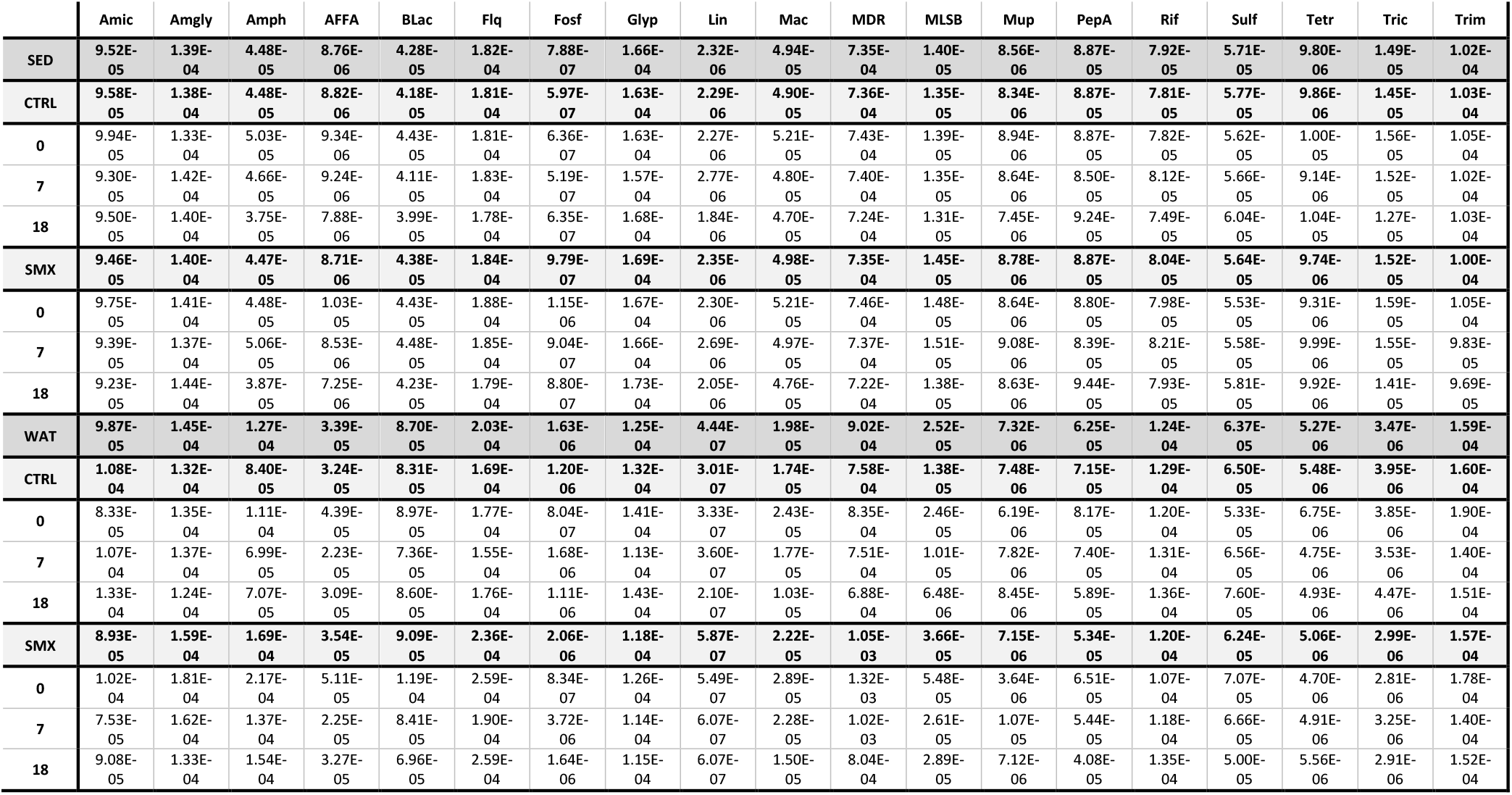
ARG relative abundance of metagenome reads according to antibiotic class. The average between three biological replicates are reported. Amic = Aminocumarin, Amgly = aminoglycoside, Amph = amphenicol, AFFA = antibacterial free fatty acids, BLac = beta-lactam, Flq = fluoroquinolone, Fosf = fosfomycin, Glyp = glycopeptide, Linc = lincosamide, Mac = macrolide, MDR = multi-drug resistance, MLSB = macrolide-lincosamide-streptogramin B resistance, Mup = mupirocin, PepA = peptide antibiotic, Rif = rifamycin, Sulf = sulfonamide, Tet = tetracycline, Tric = triclosan, Trim = trimethoprim; SED = Sediment, WAT = Water, CTRL = Control, SMX = Sulfamethoxazole

|  | Amic | Amgly | Amph | AFFA | BLac | Flq | Fosf | Glyp | Lin | Mac | MDR | MLSB | Mup | PepA | Rif | Sulf | Tetr | Tric | Trim |
| --- | --- | --- | --- | --- | --- | --- | --- | --- | --- | --- | --- | --- | --- | --- | --- | --- | --- | --- | --- |
| SED | 9.52E-05 | 1.39E-04 | 4.48E-05 | 8.76E-06 | 4.28E-05 | 1.82E-04 | 7.88E-07 | 1.66E-04 | 2.32E-06 | 4.94E-05 | 7.35E-04 | 1.40E-05 | 8.56E-06 | 8.87E-05 | 7.92E-05 | 5.71E-05 | 9.80E-06 | 1.49E-05 | 1.02E-04 |
| CTRL | 9.58E-05 | 1.38E-04 | 4.48E-05 | 8.82E-06 | 4.18E-05 | 1.81E-04 | 5.97E-07 | 1.63E-04 | 2.29E-06 | 4.90E-05 | 7.36E-04 | 1.35E-05 | 8.34E-06 | 8.87E-05 | 7.81E-05 | 5.77E-05 | 9.86E-06 | 1.45E-05 | 1.03E-04 |
| 0 | 9.94E-05 | 1.33E-04 | 5.03E-05 | 9.34E-06 | 4.43E-05 | 1.81E-04 | 6.36E-07 | 1.63E-04 | 2.27E-06 | 5.21E-05 | 7.43E-04 | 1.39E-05 | 8.94E-06 | 8.87E-05 | 7.82E-05 | 5.62E-05 | 1.00E-05 | 1.56E-05 | 1.05E-04 |
| 7 | 9.30E-05 | 1.42E-04 | 4.66E-05 | 9.24E-06 | 4.11E-05 | 1.83E-04 | 5.19E-07 | 1.57E-04 | 2.77E-06 | 4.80E-05 | 7.40E-04 | 1.35E-05 | 8.64E-06 | 8.50E-05 | 8.12E-05 | 5.66E-05 | 9.14E-06 | 1.52E-05 | 1.02E-04 |
| 18 | 9.50E-05 | 1.40E-04 | 3.75E-05 | 7.88E-06 | 3.99E-05 | 1.78E-04 | 6.35E-07 | 1.68E-04 | 1.84E-06 | 4.70E-05 | 7.24E-04 | 1.31E-05 | 7.45E-06 | 9.24E-05 | 7.49E-05 | 6.04E-05 | 1.04E-05 | 1.27E-05 | 1.03E-04 |
| SMX | 9.46E-05 | 1.40E-04 | 4.47E-05 | 8.71E-06 | 4.38E-05 | 1.84E-04 | 9.79E-07 | 1.69E-04 | 2.35E-06 | 4.98E-05 | 7.35E-04 | 1.45E-05 | 8.78E-06 | 8.87E-05 | 8.04E-05 | 5.64E-05 | 9.74E-06 | 1.52E-05 | 1.00E-04 |
| 0 | 9.75E-05 | 1.41E-04 | 4.48E-05 | 1.03E-05 | 4.43E-05 | 1.88E-04 | 1.15E-06 | 1.67E-04 | 2.30E-06 | 5.21E-05 | 7.46E-04 | 1.48E-05 | 8.64E-06 | 8.80E-05 | 7.98E-05 | 5.53E-05 | 9.31E-06 | 1.59E-05 | 1.05E-04 |
| 7 | 9.39E-05 | 1.37E-04 | 5.06E-05 | 8.53E-06 | 4.48E-05 | 1.85E-04 | 9.04E-07 | 1.66E-04 | 2.69E-06 | 4.97E-05 | 7.37E-04 | 1.51E-05 | 9.08E-06 | 8.39E-05 | 8.21E-05 | 5.58E-05 | 9.99E-06 | 1.55E-05 | 9.83E-05 |
| 18 | 9.23E-05 | 1.44E-04 | 3.87E-05 | 7.25E-06 | 4.23E-05 | 1.79E-04 | 8.80E-07 | 1.73E-04 | 2.05E-06 | 4.76E-05 | 7.22E-04 | 1.38E-05 | 8.63E-06 | 9.44E-05 | 7.93E-05 | 5.81E-05 | 9.92E-06 | 1.41E-05 | 9.69E-05 |
| WAT | 9.87E-05 | 1.45E-04 | 1.27E-04 | 3.39E-05 | 8.70E-05 | 2.03E-04 | 1.63E-06 | 1.25E-04 | 4.44E-07 | 1.98E-05 | 9.02E-04 | 2.52E-05 | 7.32E-06 | 6.25E-05 | 1.24E-04 | 6.37E-05 | 5.27E-06 | 3.47E-06 | 1.59E-04 |
| CTRL | 1.08E-04 | 1.32E-04 | 8.40E-05 | 3.24E-05 | 8.31E-05 | 1.69E-04 | 1.20E-06 | 1.32E-04 | 3.01E-07 | 1.74E-05 | 7.58E-04 | 1.38E-05 | 7.48E-06 | 7.15E-05 | 1.29E-04 | 6.50E-05 | 5.48E-06 | 3.95E-06 | 1.60E-04 |
| 0 | 8.33E-05 | 1.35E-04 | 1.11E-04 | 4.39E-05 | 8.97E-05 | 1.77E-04 | 8.04E-07 | 1.41E-04 | 3.33E-07 | 2.43E-05 | 8.35E-04 | 2.46E-05 | 6.19E-06 | 8.17E-05 | 1.20E-04 | 5.33E-05 | 6.75E-06 | 3.85E-06 | 1.90E-04 |
| 7 | 1.07E-04 | 1.37E-04 | 6.99E-05 | 2.23E-05 | 7.36E-05 | 1.55E-04 | 1.68E-06 | 1.13E-04 | 3.60E-07 | 1.77E-05 | 7.51E-04 | 1.01E-05 | 7.82E-06 | 7.40E-05 | 1.31E-04 | 6.56E-05 | 4.75E-06 | 3.53E-06 | 1.40E-04 |
| 18 | 1.33E-04 | 1.24E-04 | 7.07E-05 | 3.09E-05 | 8.60E-05 | 1.76E-04 | 1.11E-06 | 1.43E-04 | 2.10E-07 | 1.03E-05 | 6.88E-04 | 6.48E-06 | 8.45E-06 | 5.89E-05 | 1.36E-04 | 7.60E-05 | 4.93E-06 | 4.47E-06 | 1.51E-04 |
| SMX | 8.93E-05 | 1.59E-04 | 1.69E-04 | 3.54E-05 | 9.09E-05 | 2.36E-04 | 2.06E-06 | 1.18E-04 | 5.87E-07 | 2.22E-05 | 1.05E-03 | 3.66E-05 | 7.15E-06 | 5.34E-05 | 1.20E-04 | 6.24E-05 | 5.06E-06 | 2.99E-06 | 1.57E-04 |
| 0 | 1.02E-04 | 1.81E-04 | 2.17E-04 | 5.11E-05 | 1.19E-04 | 2.59E-04 | 8.34E-07 | 1.26E-04 | 5.49E-07 | 2.89E-05 | 1.32E-03 | 5.48E-05 | 3.64E-06 | 6.51E-05 | 1.07E-04 | 7.07E-05 | 4.70E-06 | 2.81E-06 | 1.78E-04 |
| 7 | 7.53E-05 | 1.62E-04 | 1.37E-04 | 2.25E-05 | 8.41E-05 | 1.90E-04 | 3.72E-06 | 1.14E-04 | 6.07E-07 | 2.28E-05 | 1.02E-03 | 2.61E-05 | 1.07E-05 | 5.44E-05 | 1.18E-04 | 6.66E-05 | 4.91E-06 | 3.25E-06 | 1.40E-04 |
| 18 | 9.08E-05 | 1.33E-04 | 1.54E-04 | 3.27E-05 | 6.96E-05 | 2.59E-04 | 1.64E-06 | 1.15E-04 | 6.07E-07 | 1.50E-05 | 8.04E-04 | 2.89E-05 | 7.12E-06 | 4.08E-05 | 1.35E-04 | 5.00E-05 | 5.56E-06 | 2.91E-06 | 1.52E-04 |

A targeted quantitative analysis of selected ARGs and MGEs by high-throughput qPCR showed that a total of 139 and 87 ARG/MGE out of 380 targets tested were positively identified in water and sediment respectively, representing a positive hit of 36.6 % and 22.9 % (Table S1). This indicated a higher diversity of ARGs in the water fraction, although the prevalence (as ARG copies/bacterial genome equivalent) was similar in the two environments (Fig 5).

**Fig 5.**
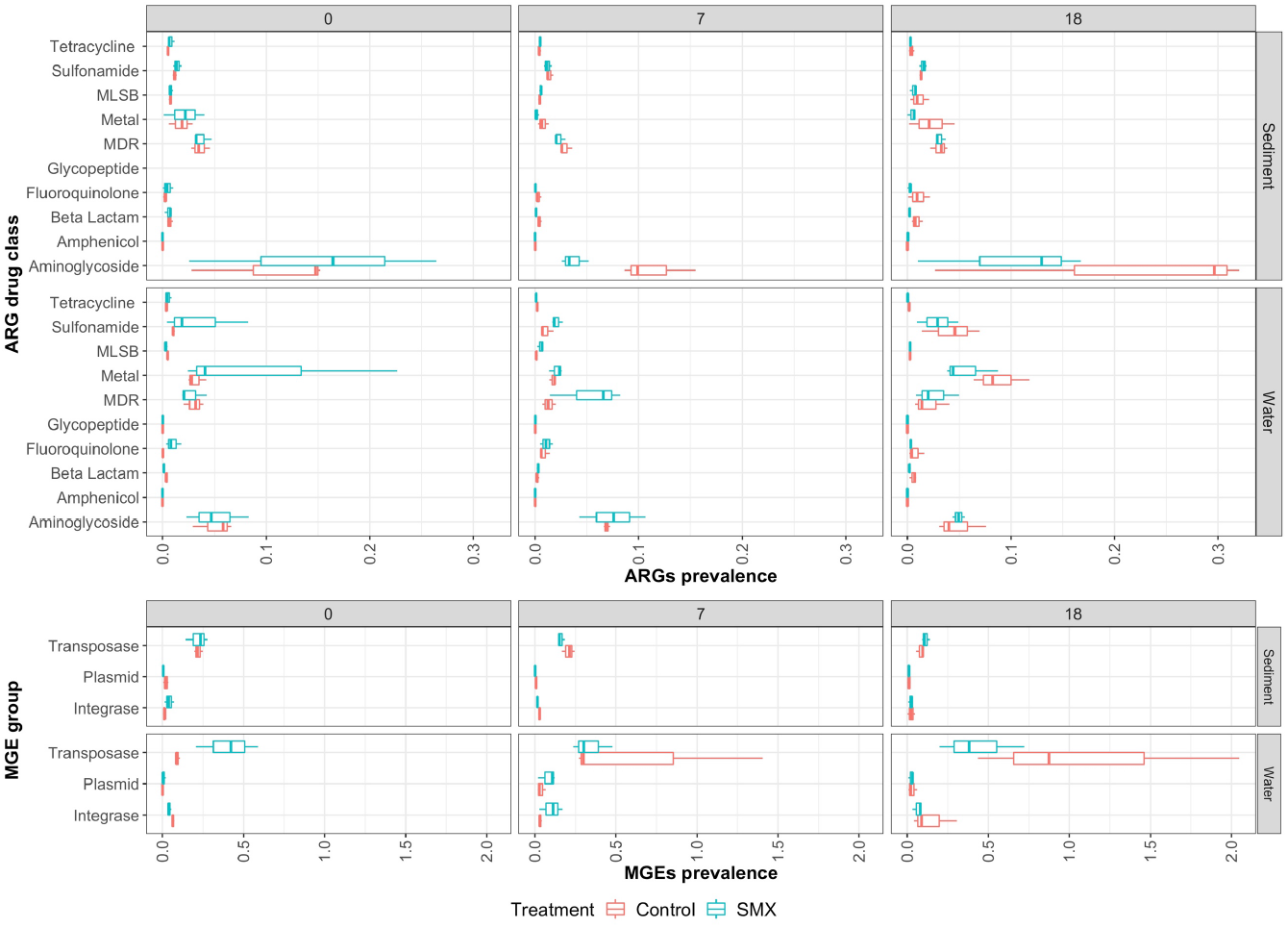
ARGs and MGEs prevalence based on high-throughput qPCR data for both water and sediment samples for the selected time point (0,7,18 days). ARG and MGE prevalence was calculated as the ratio of ARG or MGE counts/Bacterial Genome Equivalent. The median of the sum of the ratios of targets belonging to the same ARG drug class or MGE group are reported. Three biological replicates were used for each condition (Control/SMX).

The qPCR data showed significant ARG prevalence differences between the two environments (Adonis, R2=0.24, p=0.001). However, no significant changes were observed that could be explained by either the presence of SMX (Adonis water R2=0.05, p>0.05; sediment R2=0.02, p>0.05) or time (Adonis water R2=0.06, p>0.05; sediment R2=0.13, p>0.05). These results implied that the presence of SMX at the environmental concentration of <4 μg L^−1^ did not significantly affect the ARG and MGE composition of the microbiome (Fig 5).

Due to conflicting results from metagenomics and qPCR analysis for the ARG response in the water microbiome in presence of the SMX, correlation analysis of ARG prevalence detected by both methods (metagenomics and qPCR) were explored in order to define the effect of SMX on the resistome (Fig 6). Although correlations were not significant for the common antibiotic classes detected by each method individually (Fig S9), the combination of both datasets enabled identification of positive correlations between MDR, MLSB, fluoroquinolone and amphenicol ARG classes with presence of SMX in the water microbiome. Some of these classes (MDR and fluoroquinolone) also had a strong positive correlation with MGE such as plasmids and integrases which did not themselves show a significant correlation to SMX. This suggested that the presence of SMX did not affect the spreading of ARG, even though an enrichment of selected classes of ARG might have occurred in the water microbiome (Fig 6).

**Fig 6.**
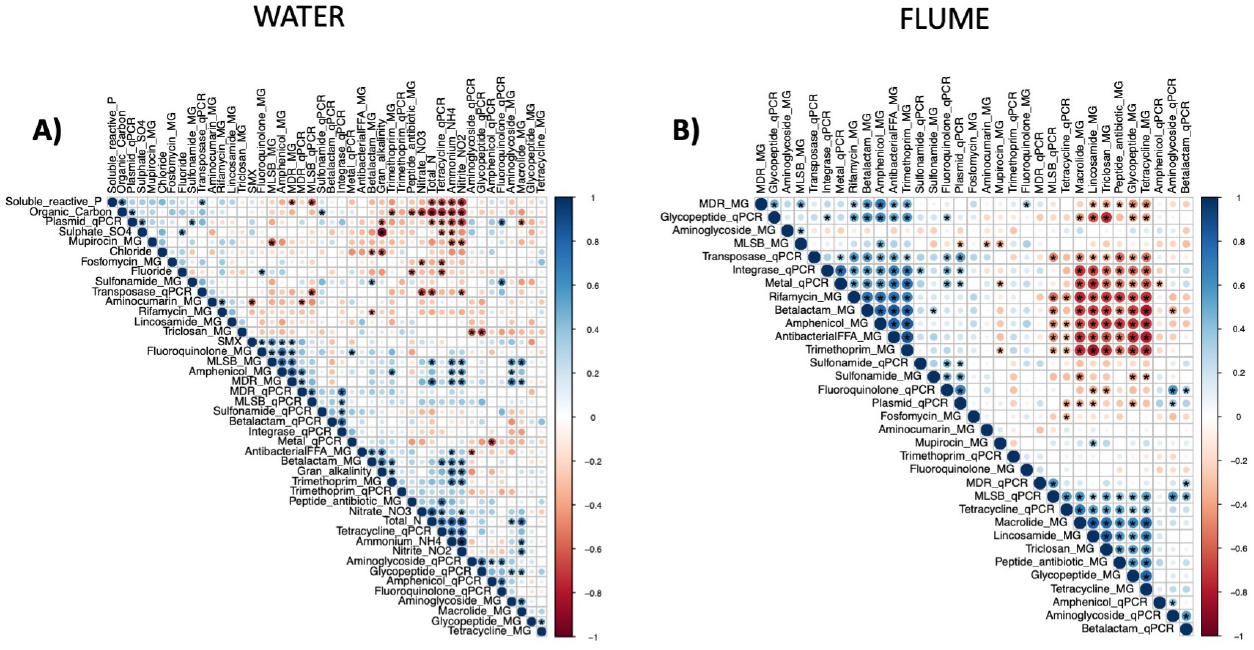
Spearman rank correlation between ARG prevalence of classes of antibiotic resistance detected by metagenome (MG) and qPCR analysis. The symbol (*) represents significant correlations (p<0.05), circle size corresponds to the correlation value, while the colour to either positive (blue) or negative (red) correlation. A) Water. The nutrient chemical data available for the water microbiome were also included in the analysis. B) Flume (water+sediment).

The taxonomic distribution of the ARG reads detected in the metagenomes showed that the majority were assigned to the Proteobacteria phylum suggesting that this phylum is the greatest reservoir for ARGs in the river environment (Fig S10). Within the 812 MAGs recovered from the water microbiome (4 Archaea and 808 Bacteria) (Fig S11), 22 presented ARGs. Amongst these MAGs, 14/22 were assigned to Proteobacteria including potentially pathogenic genera such as *Pseudomonas, Shewanella, Acinetobacter, Legionella* and environmental relevant genera such as *Methyloversatilis, Caedimonas, Arenimonas* and *Zoogloea*. The remaining 8 MAGs containing ARGs were assigned to the Actinobacteriota genera *Mycolicibacterium, Micropruina* and *Mycobacterium*, and the Chloroflexota and Verrucomicrobiota families of UBA6265 and Opitutaceae respectively (Table S2). For 9/22 of these MAGs multiple ARGs were identified providing resistance to multiple drugs such as for the MAG related to the human pathogen *Acinetobacter beijerinckii* which showed 9 different genes related to MDR efflux pumps and tetracycline resistance (Table S2).

Although the prevalence of ARGs was low in both microbiomes, expression of antimicrobial resistance proteins (ARPs) was investigated through recovery of metaproteomes. Sediment metaproteomes proved difficult to recover due to the nature of the sample and co-extraction of impurities that could not be separated from the samples; thus, metaproteomic data were not collected for the sediment microbiome. Metaproteomic analysis of the water fraction showed that it was possible to detect proteins involved in the resistance to rifampicin, elfamycin, kirromycin, GE2270A and daptomycin, some more evenly detected than others (Fig 7C). It is likely that all these ARPs are chromosomally encoded. However, no significant difference in their expression was observed between the control and the SMX group (t-test p<0.001) suggesting that their expression is not linked or influenced by environmental concentration of SMX (Fig. 7A and 7B).

**Fig 7.**
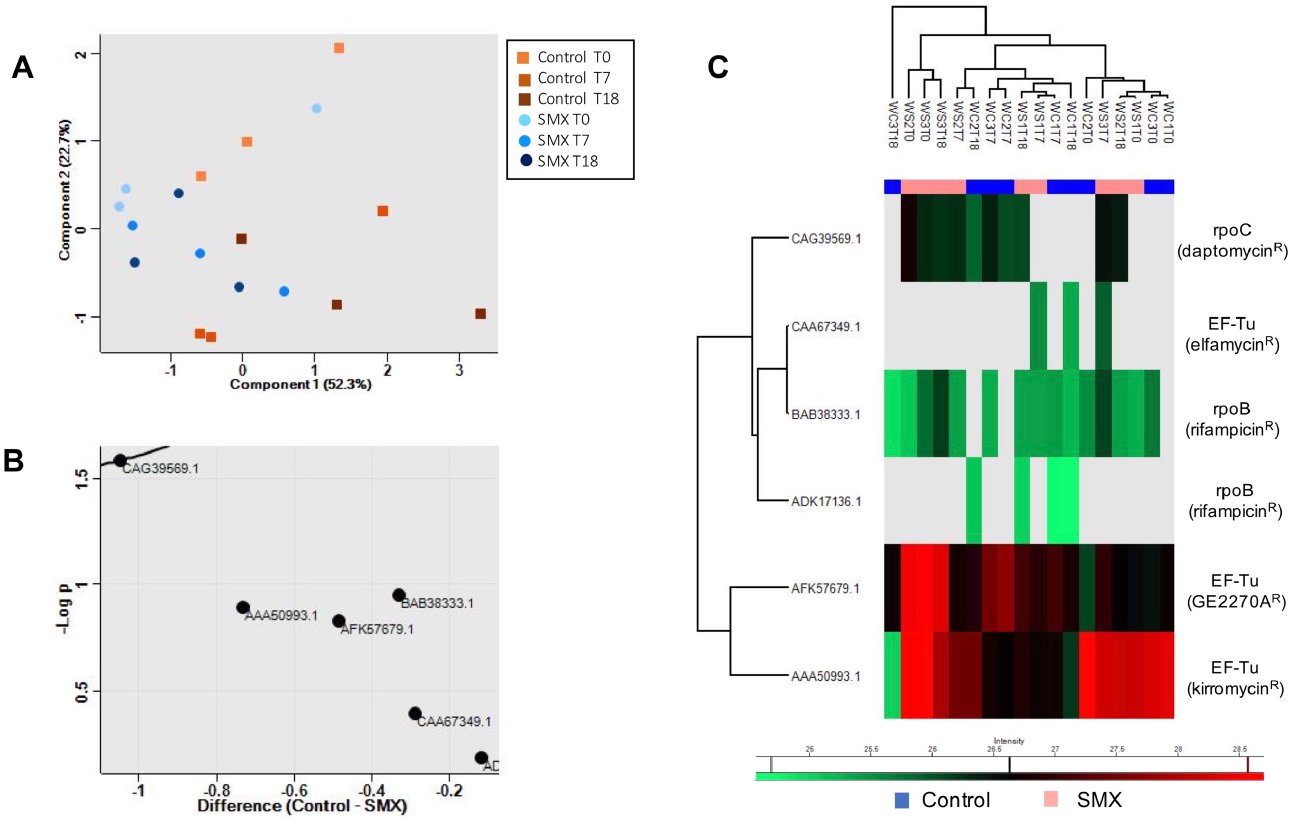
Antimicrobial resistant proteins (ARPs) detected in the water metaproteomic samples. A) Samples separation by PCA coloured according to time point and treatment (control vs SMX). B) Volcano plot representing the difference in the log_2_ label-free quantitative (LFQ) intensity values of the detected ARPs between control and SMX samples and the log probability of the observed differences. The line represents the threshold for the significant p-value. C) Heat-map of the log_2_ LFQ intensity of the detected ARPs in each individual sample. Grey colour = NaN.

## Discussion

The replication of freshwater environments such as rivers in the laboratory is difficult and microcosms studies were usually performed on static or shaking systems which only partially resembled the natural environment dynamic characteristics (Grenni et al., 2019; Patrolecco et al., 2018; Tong and Xie, 2019; Vila-Costa et al., 2017). In the current study flumes were set up in closed *in vitro* systems where water flowed on a sediment bed to better mimic river and environmental conditions. The use of a controlled closed system allowed the reduction of impact from different variables such as continuous addition of nutrients, xenobiotics and diverse microbiomes to the system. We were therefore able to test the effects of SMX addition on the riverine microbiome in both the sediment and the water column in semi-natural conditions. This system has proved to be robust and reliable with limited variation between flumes.

The introduction of an environmentally relevant concentration of SMX (<4 μg L^−1^) did not significantly affect the flume sediment microbiome composition; nor did it affect the ARG diversity or prevalence. Conversely, a limited response was observed in the water microbiome, showing a correlation between the SMX presence with MDR and fluoroquinolone resistance classes but not with sulfonamide resistance ARGs as originally hypothesized. The ARG classes that did correlate were linked to MGEs, which however did not show a significant correlation with the presence of SMX. These results suggest that the release of low levels of SMX (<4 μg L^−1^) from wastewater effluents or other sources into the river may have a limited impact on ARG selection in the riverine microbiome without necessarily promoting ARG spread during short periods of exposure as tested in this study. However, longer periods of continuous exposure to low concentrations of antibiotics could present completely different outcomes that shape the microbial community resistome. A recent river water microcosm study monitored selected ARG sulfonamide resistance targets (*sul1* and *sul2*) in the microbial community of a biofilm that formed on pebbles in the presence of 1 μg L^−1^ and 5 μg L^−1^ SMX over 60 days. ARG prevalence at a low concentration of SMX (<1 μg L^−1^) was not affected in the first 30 days of exposure but an increase of ARG was observed in the following 30 days. This led to a hypothesis that adaptation of the indigenous community to degrade the compound was mainly involved in the first half of the experiment thereby providing resistance to SMX, while the spread of ARGs was promoted in the second half (Vila-Costa et al., 2017). Our study showed an impact of SMX on ARGs in the water column as opposed to the sediment and no significant impact on the community diversity in either the water or the sediment fractions.

This study is also the first report of ARP prevalence in a natural riverine environment. It was interesting to note that in the protein fraction analysed, no DHSP enzymes resistant to sulfonamide were detected, confuting our hypothesis that the presence of SMX would promote the expression of more proteins related to its resistance in the community. The proteomic results showed that although SMX might have an effect on the microbiome composition at a genetic level, the expression of ARGs was not significantly promoted in these conditions, although differential sensitivity of these techniques makes comparisons difficult.

Comparison analysis with preliminary results from a pilot study conducted with samples collected in May 2018 (supplementary material Table S3 and Fig S12 - S14) showed a potential effect due to seasonal variability of the starting microbiome used in the flumes experiment. The pilot study showed an increase in ARGs related to sulfonamide resistant and MGEs after exposure to sub-lethal concentration of SMX. However, the insufficient high-throughput qPCR data available for this pilot study did not allow us to draw conclusive interpretations in relation to ARGs spreading in the conditions tested. The main experiment conducted in 2019 was an attempt to replicate the same conditions but seasonal variation of the microbial community was observed between the studies which may explain the differential impact of SMX on the microbial community. This suggests that further studies are needed to assess the effect of sublethal concentrations of antimicrobials in natural environmental microbiomes collected in different seasons and exposed to different environmental conditions as these might have an impact on their response. However, it is clear that there is considerable variation in the estimation of PNEC for selection of resistance to sulfonamides (Mortimer et al., 2020), which supports the variation between our pilot study and the study reported here in detail. We believe that estimating PNEC for environmental microbiomes is extremally challenging and will be subject to considerable variation depending on the large number of variables that determine microbiome functions.

## Conclusions

This *in vitro* flume system proved to be a reliable semi-natural system to implement further studies on the effects of release of xenobiotics in freshwater natural environments such as rivers. Microcosms integrating semi-realistic features of rivers will be necessary in further studies to test longer exposure effects to sub-lethal concentrations of individual xenobiotics. Complex combinations of pollutants exist in effluents and this will need to be simulated in future studies. Integration of these *in vitro* experiments with monitoring and surveillance studies will help to better predict AMR levels in the environment and potential risk to human health, providing support for mitigation strategies.

## Supplementary files

**Additional file 1:** Supplementary methods and results.

**Table S1:** MAGs containing ARGs

## Availability of data and materials

The datasets generated and analysed during the current study are available in the Short Read Archive (Bioproject PRJNA693684). The mass spectrometry proteomic data have been deposited to the ProteomeXchange Consortium via the PRIDE partner repository with the dataset identifier PXD023822.

## Acknowledgments

This work was supported by the Natural Environment Research Council (grant numbers NE/N019857/1, NE/N019687/1, NE/S013539/1, NE/S008721/1); the Medical Research Council (UKRI MRC) Korean Partnering Scheme (grant number MC_PC_18014); the Korea Health Technology R&D Project through the Korea Health Industry Development Institute (KHIDI), funded by the Ministry of Health & Welfare, Republic of Korea (grant number: HI18C2063).

The authors would also like to acknowledge the help received from: Dr. Cleidiane Zampronio at WPH Proteomics RTP, University of Warwick - metaproteomic samples preparation; Ian Baylis at the School of Engineering, University of Warwick - maintenance of the flumes system; Andrew Murphy, James Delaney, Valentin Waschulin and Dr. Ian Lidbury, School of Life Sciences, University of Warwick - river sampling; Dr. Mike Bowes’s lab at the UK Centre for Ecology and Hydrology - nutrient chemical analysis under the UKRI service scheme.

