## Additional file 1 for "Impact of sulfamethoxazole on a riverine microbiome"

**Nutrients and SMX chemical analysis in flumes**

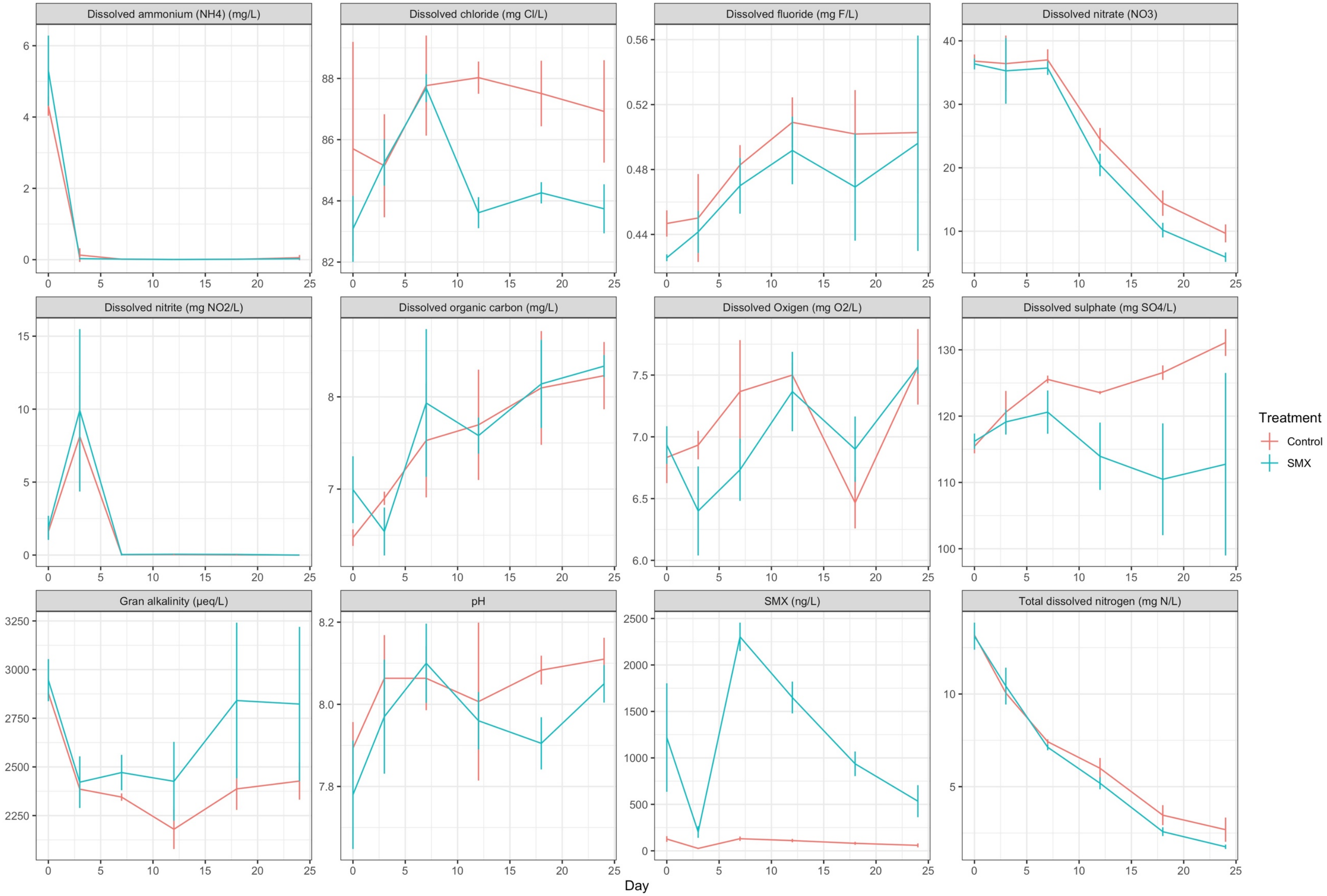


Fig S1. Nutrient chemicals concentrations in water in both groups (Control and SMX). Average concentration of three biological replicates and standard deviations are reported for each time point.

### Alpha diversity

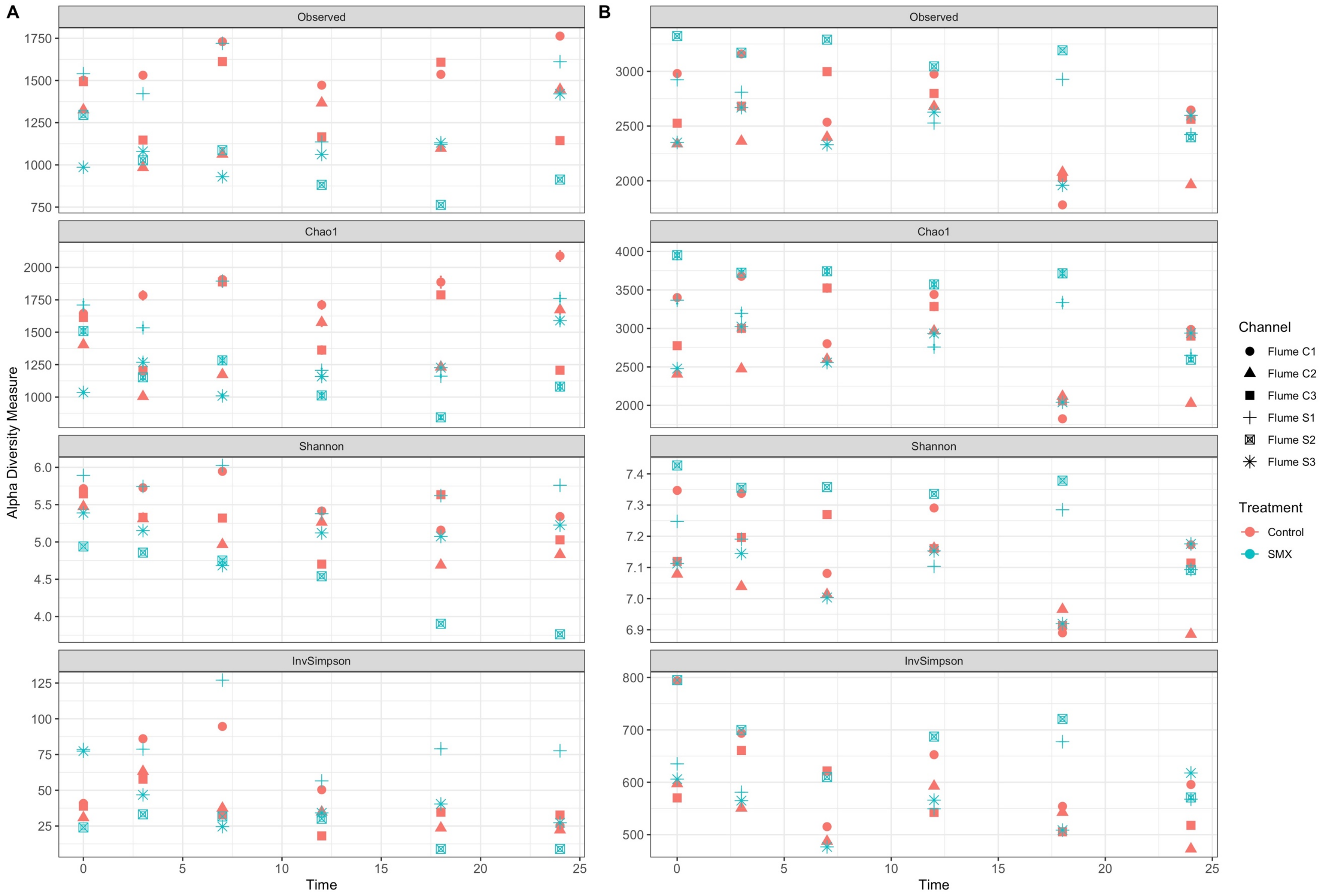


Fig S2. Alpha diversity of microbial communities in water (A) and sediment (B). Observed species, Chao1, Shannon and InvSimpson indices were calculated for each sample based on 16S rRNA gene amplicon datasets.

### Microbiome structure

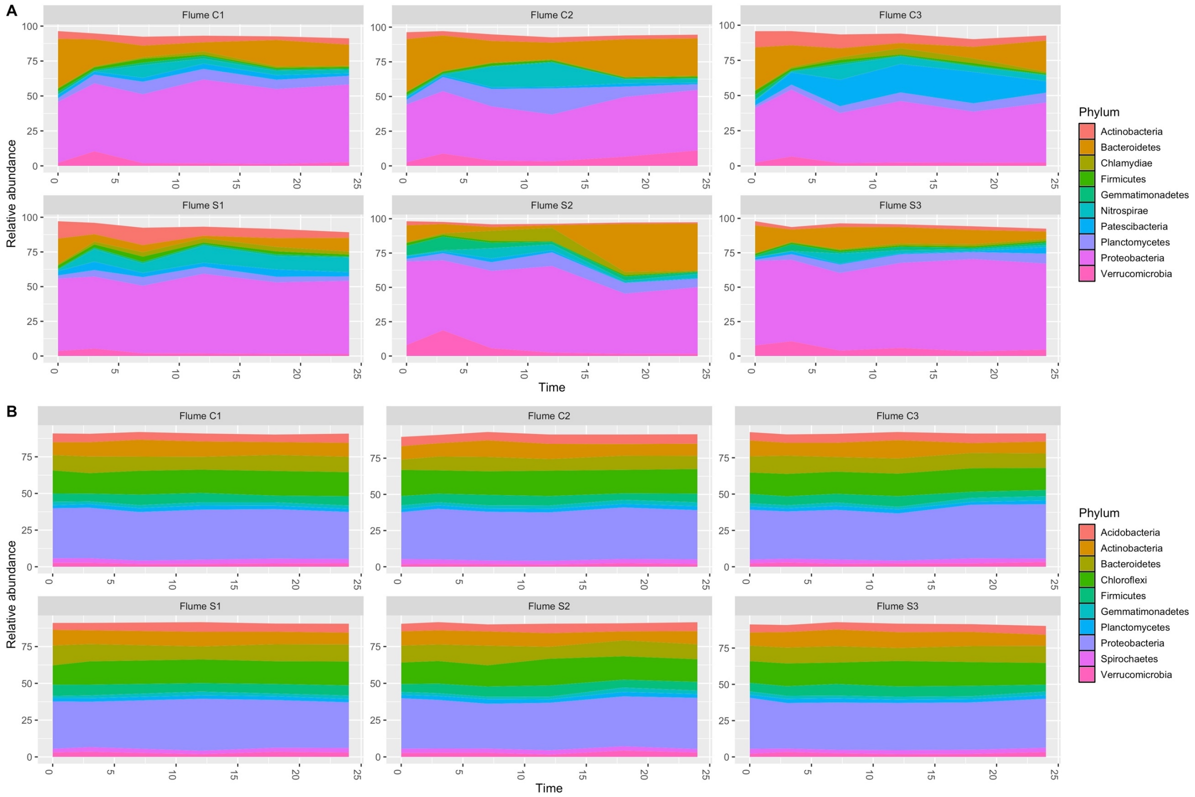


Fig S3. Taxonomic characterization of microbial communities in water (A) and sediment (B). 16S rRNA gene amplicon relative abundance of top 10 Phyla is represented for each flume overtime (0-24 days). Flumes C1-C3: Control; Flume S1-S3: amended with Sulfamethoxazole (SMX).

### Beta diversity

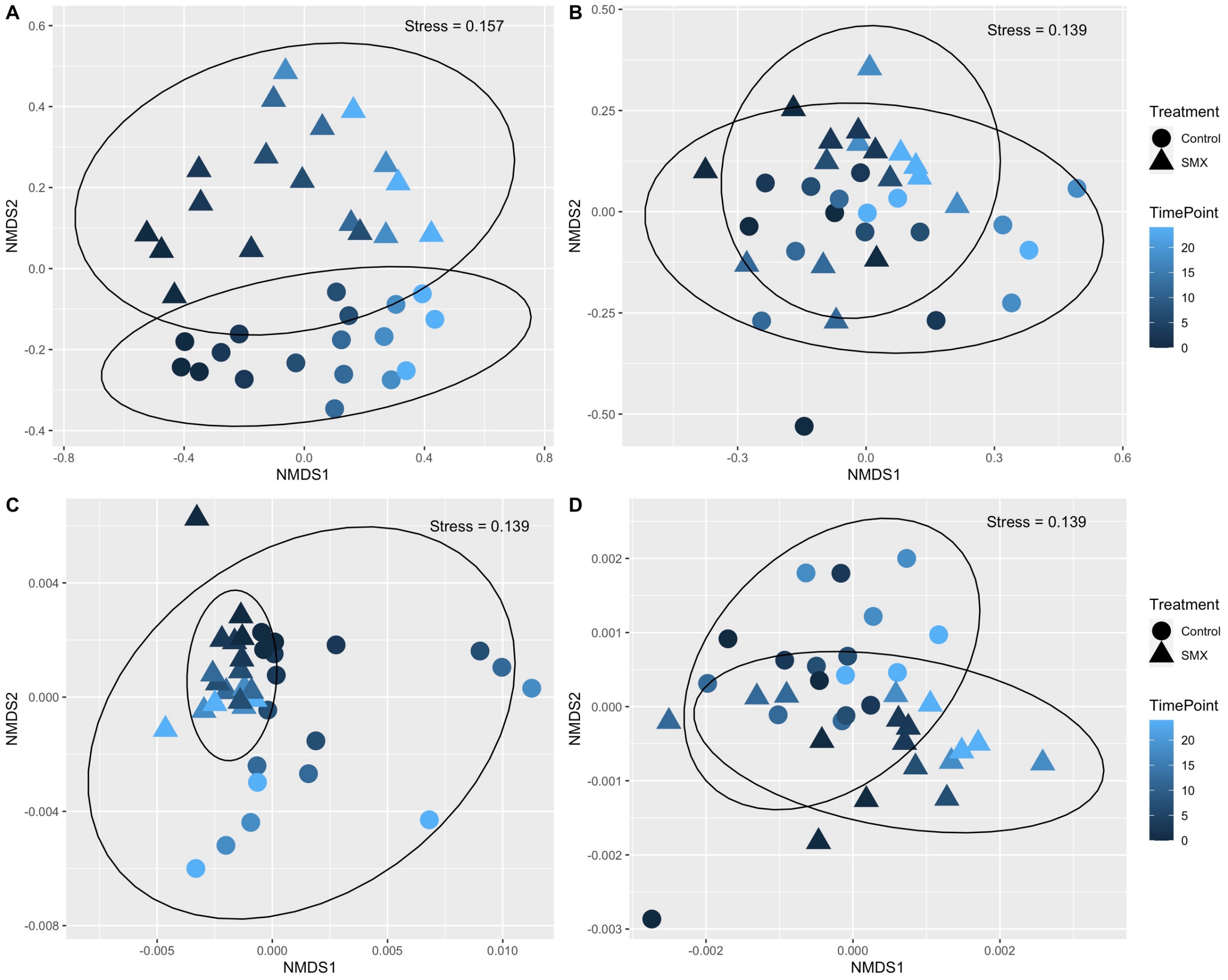


Fig S4. Microbial community ordination by NMDS of Jaccard and weighted UniFrac distances based on 16S rRNA gene amplicon datasets. A) Water - Jaccard; B) Sediment - Jaccard; C) Water – Weighted UniFrac; D) Sediment – Weighted UniFrac. Ellipses represent sample grouping by treatment (Control-SMX).

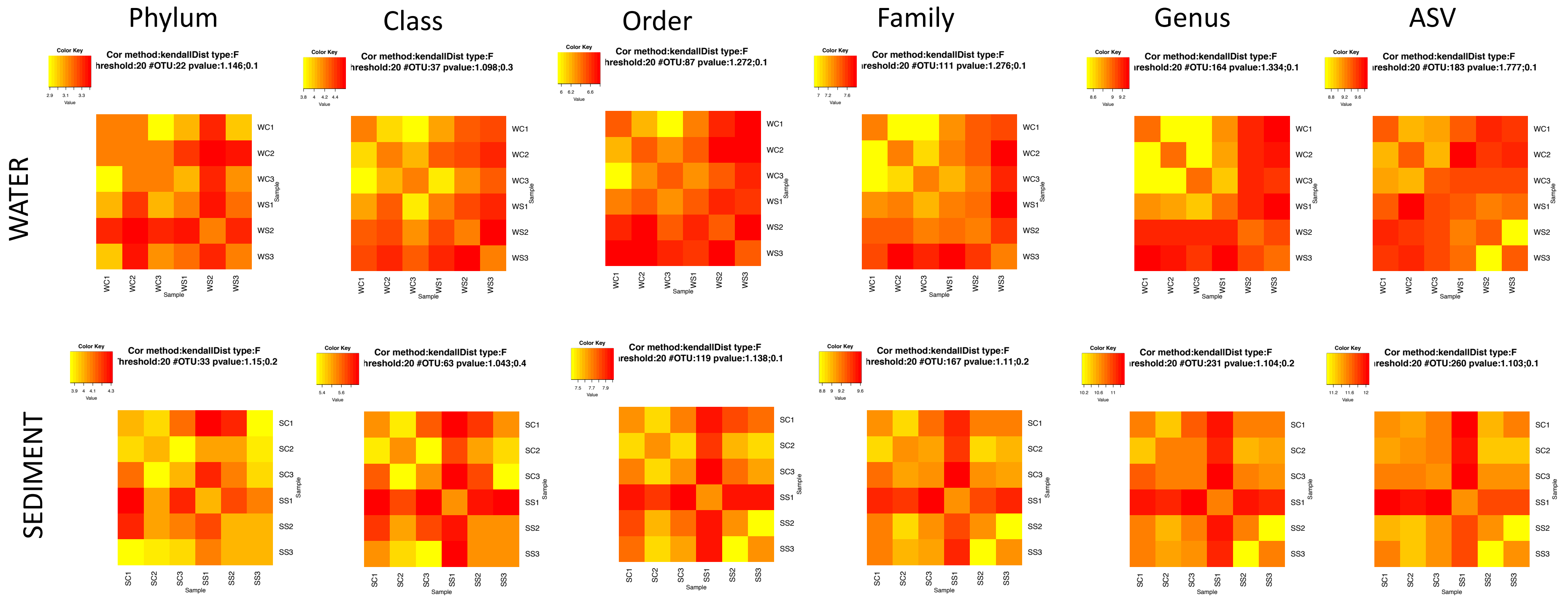


Fig S5. Nonparametric microbial interdependence temporal correlation profile at different taxonomic levels and ASV level (based on 16S rRNA gene amplicon datasets).

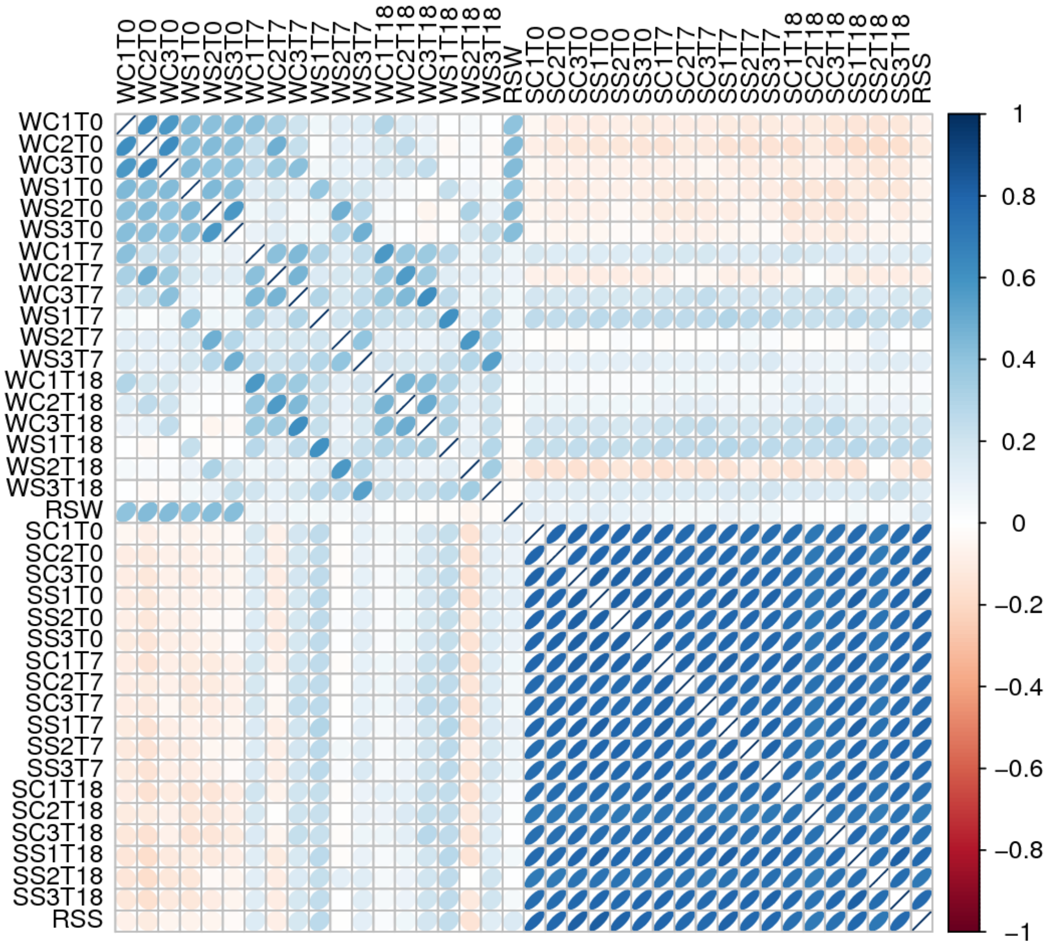


Fig S6. Correlation plot between the sequenced metagenomes for sediment and water (time points 0, 7 and 18 days). Blue represents a positive correlation, while red a negative correlation. The deeper the colour is, the greater the association.

Strong positive correlations were observed between the time point 0 days of samples collected from the flume system and the original samples collected from the river, suggesting that the starting microbiome in the flume system was a good representation of the natural microbiome present in the river.

### Metagenome functional analysis

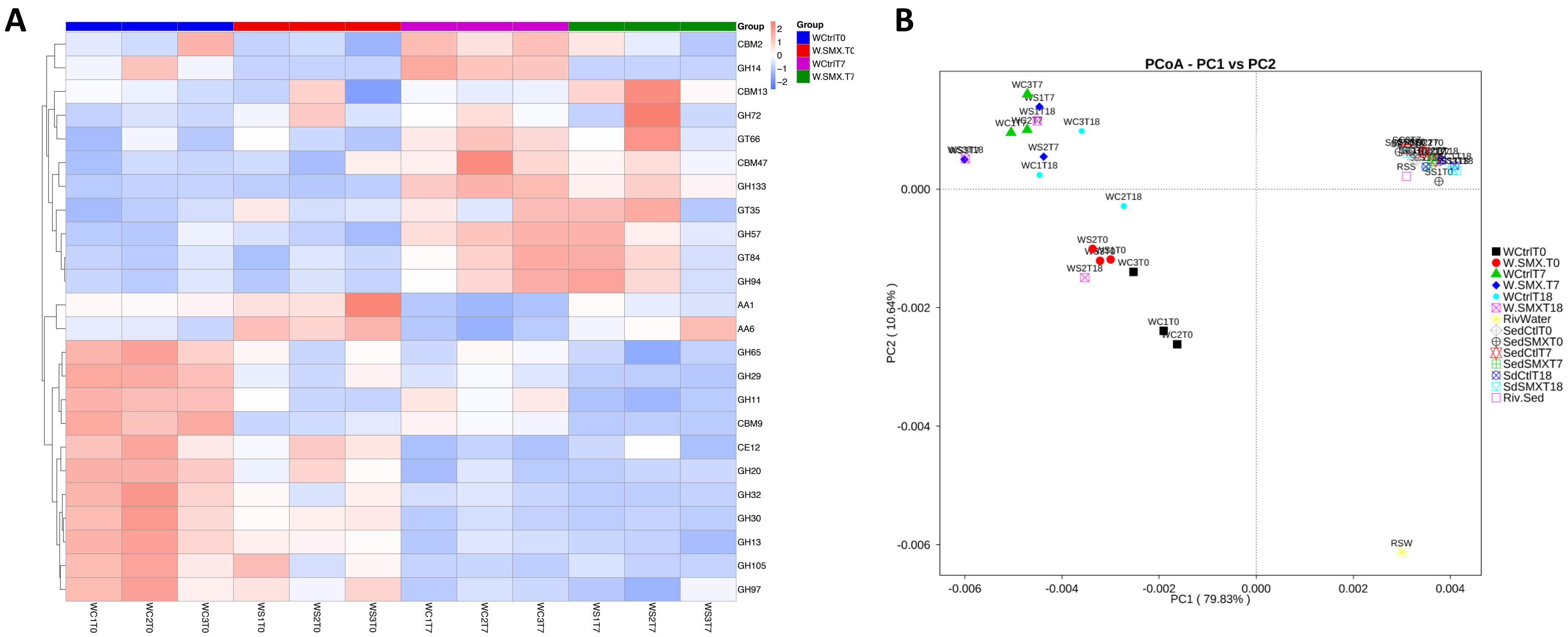


Fig S7. Metagenomic functional annotation based on CAZy database. A) Biomarker identification in water samples between Control and SMX groups at time point 0 and 7 days. B) Principal coordinates analysis based on Bray-Curtis distance of metagenomes carbohydrate functional annotation of all samples.

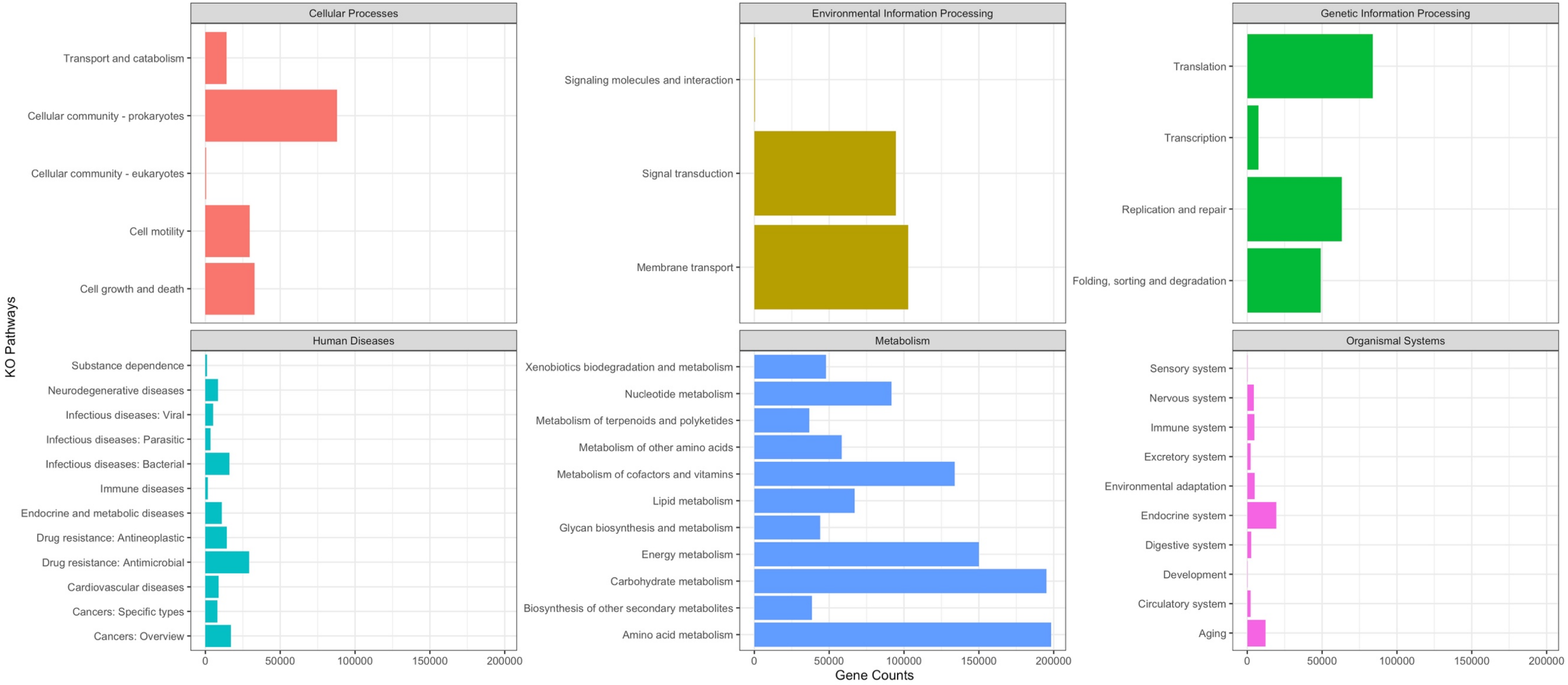


Fig S8. Total number of genes identified in all metagenomes for each KEGG pathway.

Table S1. High-throughput qPCR primer counts of positive detection according to drug class resistance classification of targets tested.

| Target class | Primer target | Total tested | Water positive | Sediment positive |
|---|---|---|---|---|
| ARG | **Aminoglycoside** | 61 | 23 | 15 |
| | **Amphenicol** | 17 | 4 | 2 |
| | **Beta Lactam** | 55 | 16 | 6 |
| | **Fluoroquinolone** | 12 | 4 | 2 |
| | **Glycopeptide** | 24 | 5 | 0 |
| | **MDR** | 35 | 13 | 8 |
| | **Metal** | 10 | 6 | 4 |
| | **MLSB** | 44 | 15 | 13 |
| | **Sulfonamide** | 4 | 2 | 2 |
| | **Tetracycline** | 28 | 15 | 13 |
| | **Trimethoprim** | 20 | 3 | 0 |
| MGE | **Integrase** | 4 | 3 | 3 |
| | **Plasmid** | 17 | 5 | 4 |
| | **Transposase** | 35 | 22 | 14 |
| | **Other** | 14 | 3 | 1 |
| **Grand Total** | | **380** | **139** | **87** |

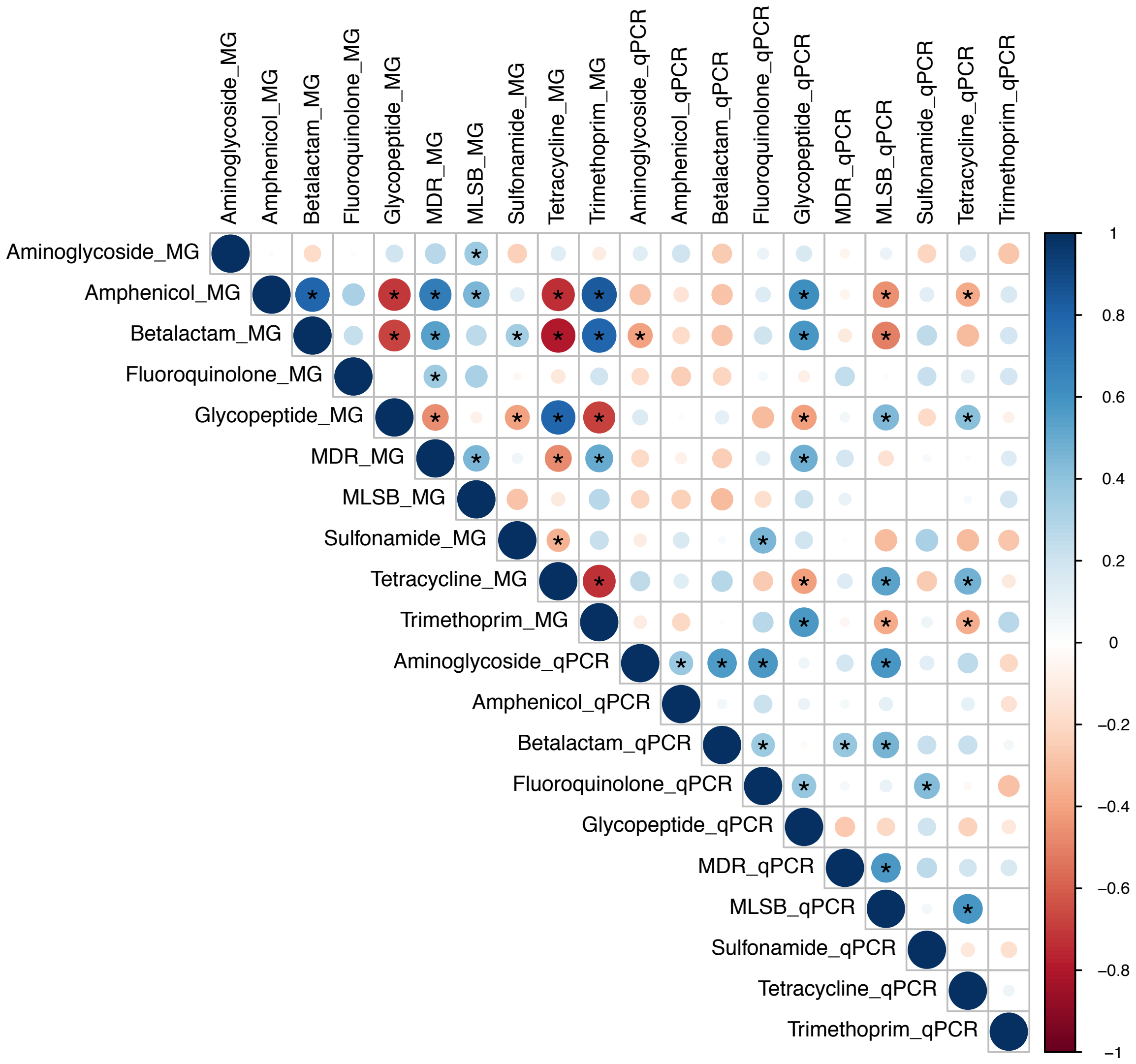


Fig S9. Spearman rank correlation of common antibiotic resistance classes of ARGs detected by metagenome and qPCR analysis. The symbol (*) represents significant correlations ($p<0.05$).

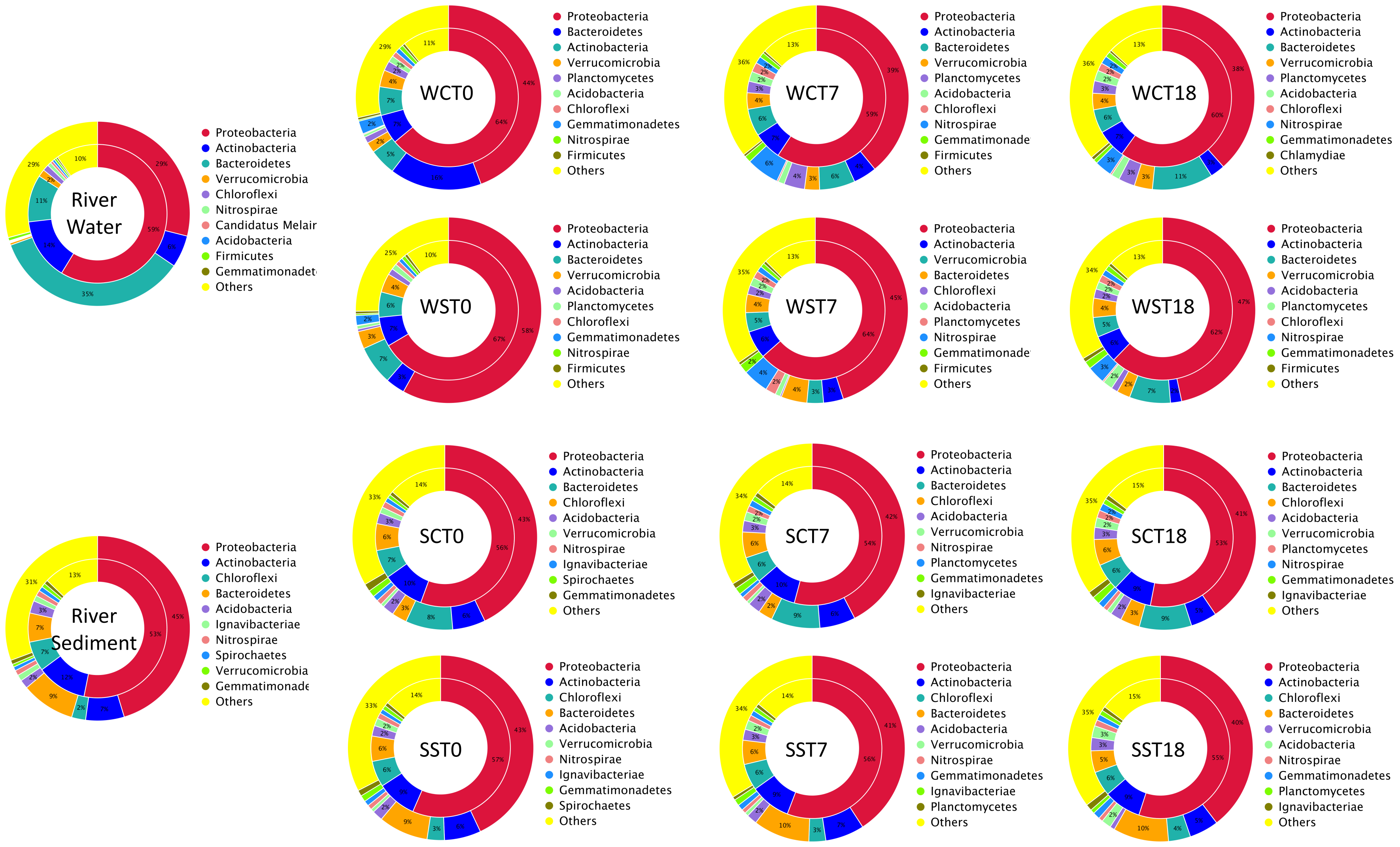


Fig S10. Proportion of different phyla in all genes and ARGs in each metagenome. Inner ring = proportion of different phyla contained ARGs; Outer ring = proportion of genes in different phyla of this group.

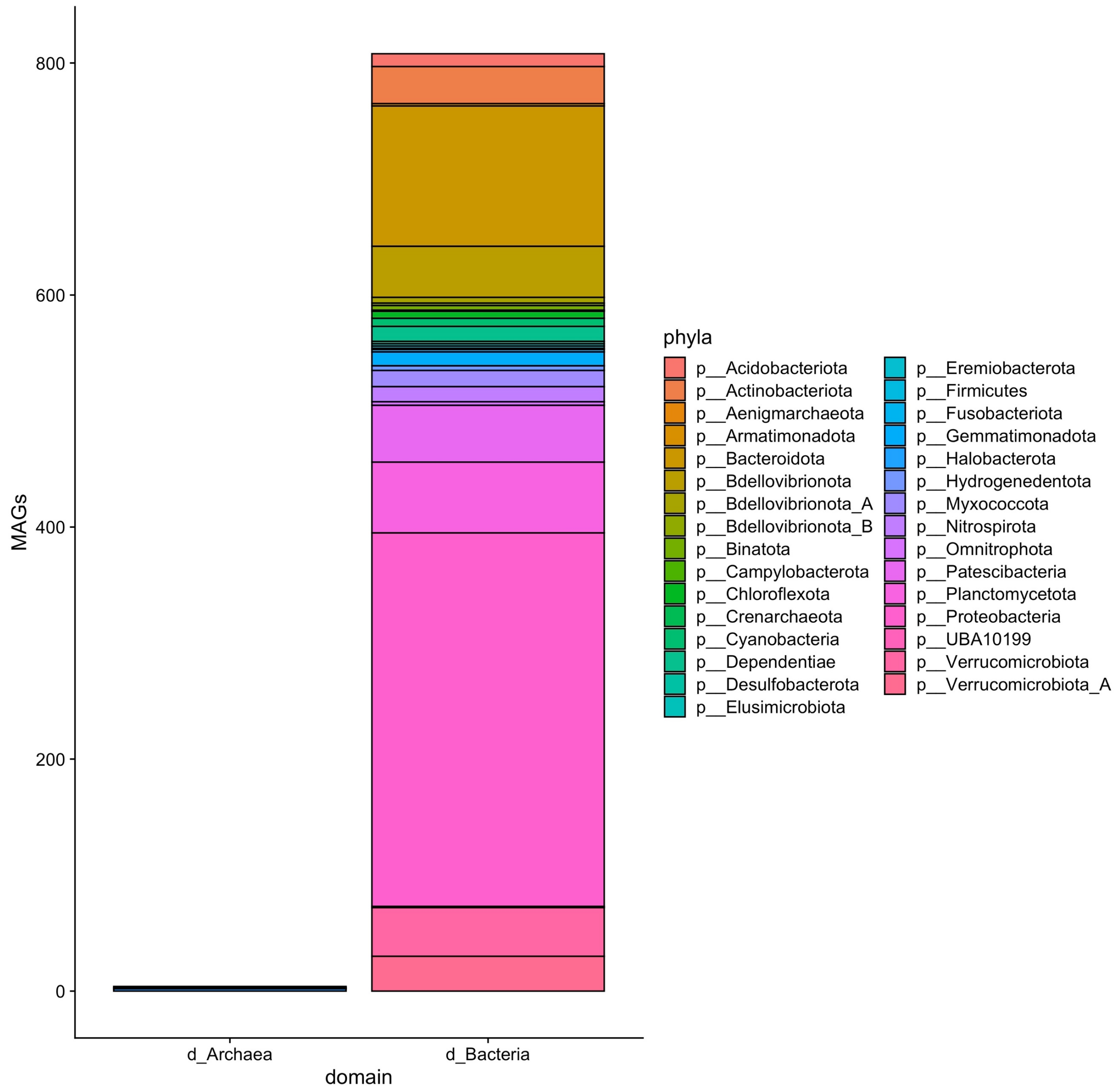


Fig S11. Taxonomic distribution (at phylum level) of all MAGs recovered from the water metagenomes.

**Pilot study (May 2018)**

**Methods**

River water and sediment were collected from the River Sowe, UK in May 2018 and immediately used to set up flumes as described in the main manuscript. Method of addition and concentration of SMX added were the same as in the main study performed in April 2019. Sampling was done at the same intervals for both experiments (0,3,7,12,18,24 days) for both water and sediment. However, in the pilot study smaller amounts of samples were collected in comparison to the 2019 experiment. In the pilot study, 0.1 L of water were filtered (0.22 μm PES filter) and 0.5 g sediment were used for DNA extraction for 16S rRNA gene V3-V4 amplicon sequencing (all samples) and high-throughput qPCR. Only a limited selection of samples (water and sediment DNA at time points 0 and 18 days for only one flume per treatment- control and SMX) were analyzed by high-throughput qPCR due to limited DNA quantities recovered. 16S rRNA gene amplicon sequences were processed using qiime2 and R using the same DADA filtering parameters used to process data from 2019. Rarefaction of the combined 2018 and 2019 dataset was performed at 12000 sequences per samples. Statistical analyses were also performed in R.

Chemical analyses of nutrients and SMX were performed on samples collected from individual flumes and pulled according to the treatment, due to the volumes of water required for some of the analysis.

**Results**

Table S3. Weather average conditions during the months of sampling for the pilot and main studies in Stoneleigh, UK where samples were collected from the River Sowe. Data were retrieved from https://www.timeanddate.com/weather/ on 23/10/2020

| Study | Month - Year | Temperature (°C) | Humidity (%) | Pressure (mbar) | Sample collection date |
|---|---|---|---|---|---|
| Pilot | May 2018 | Max 25 (on 07/05)<br>Min 2 (on 01/05)<br>Average 14 | Max 100 (on 02/05)<br>Min 31 (on 14/05)<br>Average 74 | Max 1030 (on 02/05)<br>Min 998 (on 02/05)<br>Average 1018 | 22/05/2018 |
| Main | April 2019 | Max 22 (on 19/04)<br>Min -1 (on 14/04)<br>Average 9 | Max 100 (on 03/04)<br>Min 29 (on 20/04)<br>Average 74 | Max 1032 (on 03/04)<br>Min 990 (on 04/04)<br>Average 1014 | 17/04/2019 |

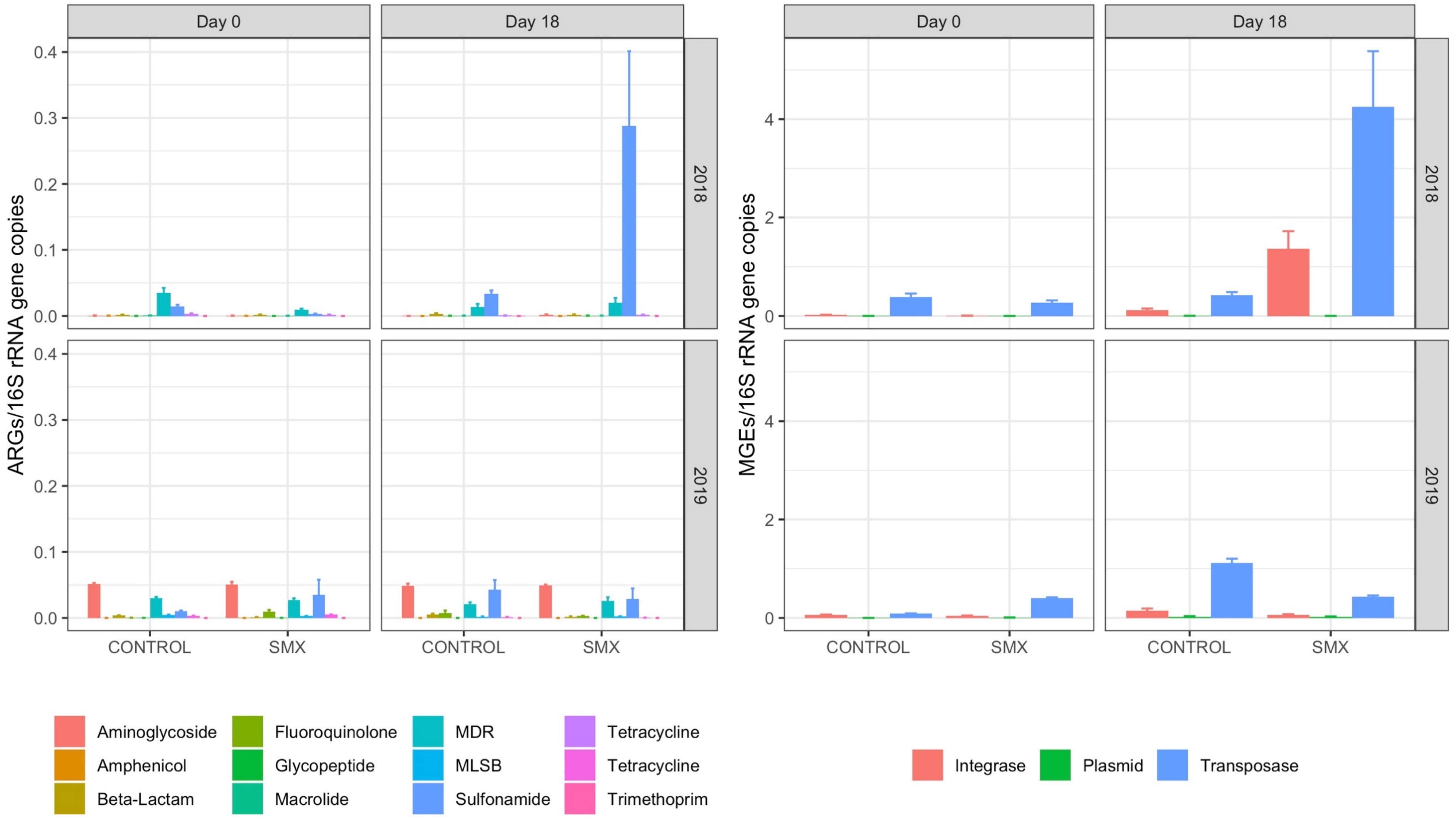


Fig S12. ARG detected by high-throughput qPCR in the samples collected in the pilot study (year 2018) and the main experiment (year 2019). For the pilot study only 1 sample per treatment (Control vs SMX) at time point 0 and 18 days were used in the analysis due to limited DNA recovered. Triplicate for each treatment at both time points (0 and 18 days) were available for analysis for the main experiment (year 2019).

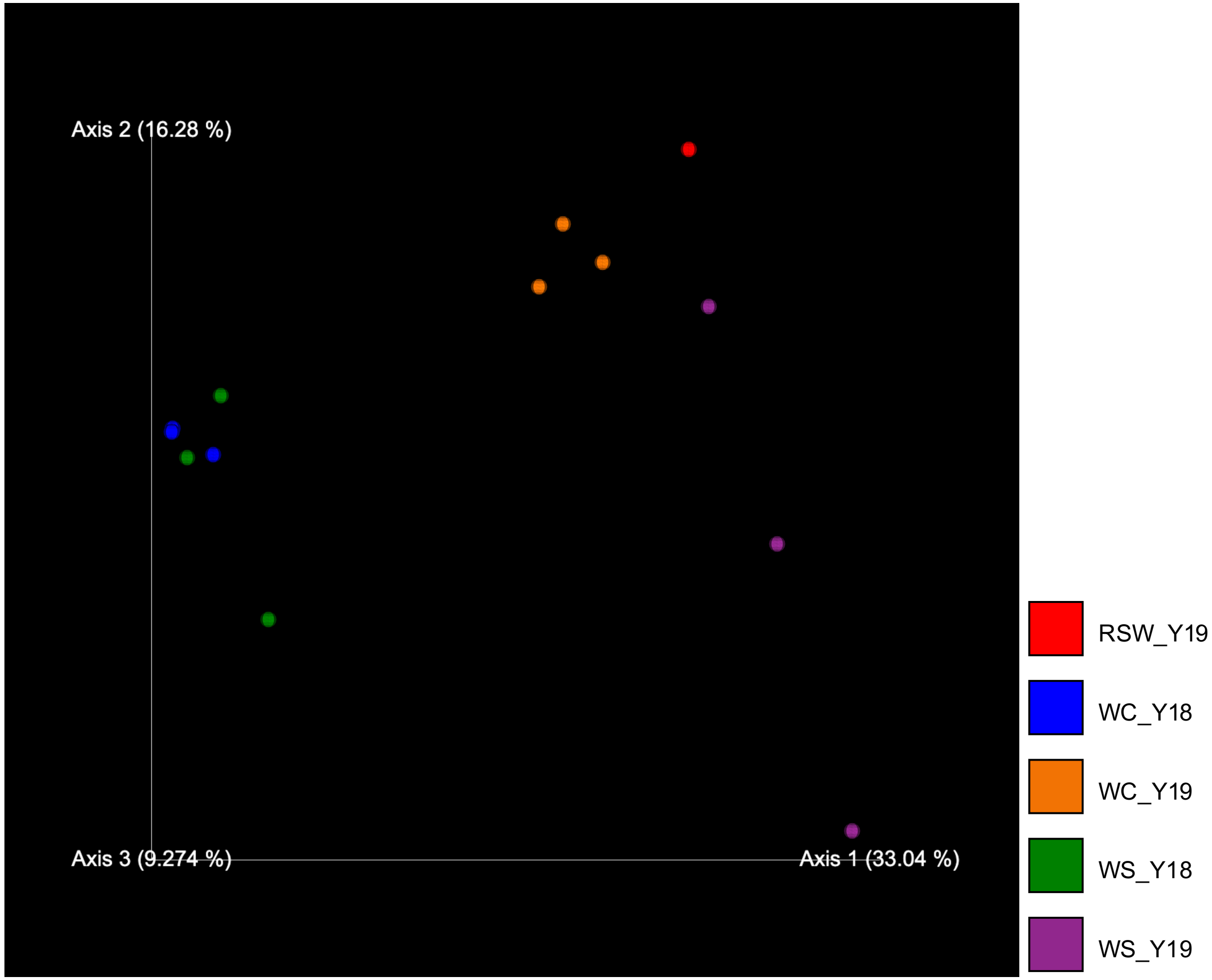


Fig S13. Comparison of water samples 16S rRNA gene diversity by Bray-Curtis PCoA between the pilot study and the main study at time point 0 days. RSW_Y19: River Sowe water (year 2019); Flume samples: WC_Y18: water control year 2018, WC_Y19: water control year 2019, WS_Y18: water + SMX year 2018, WS_Y19: water + SMX year 2019.

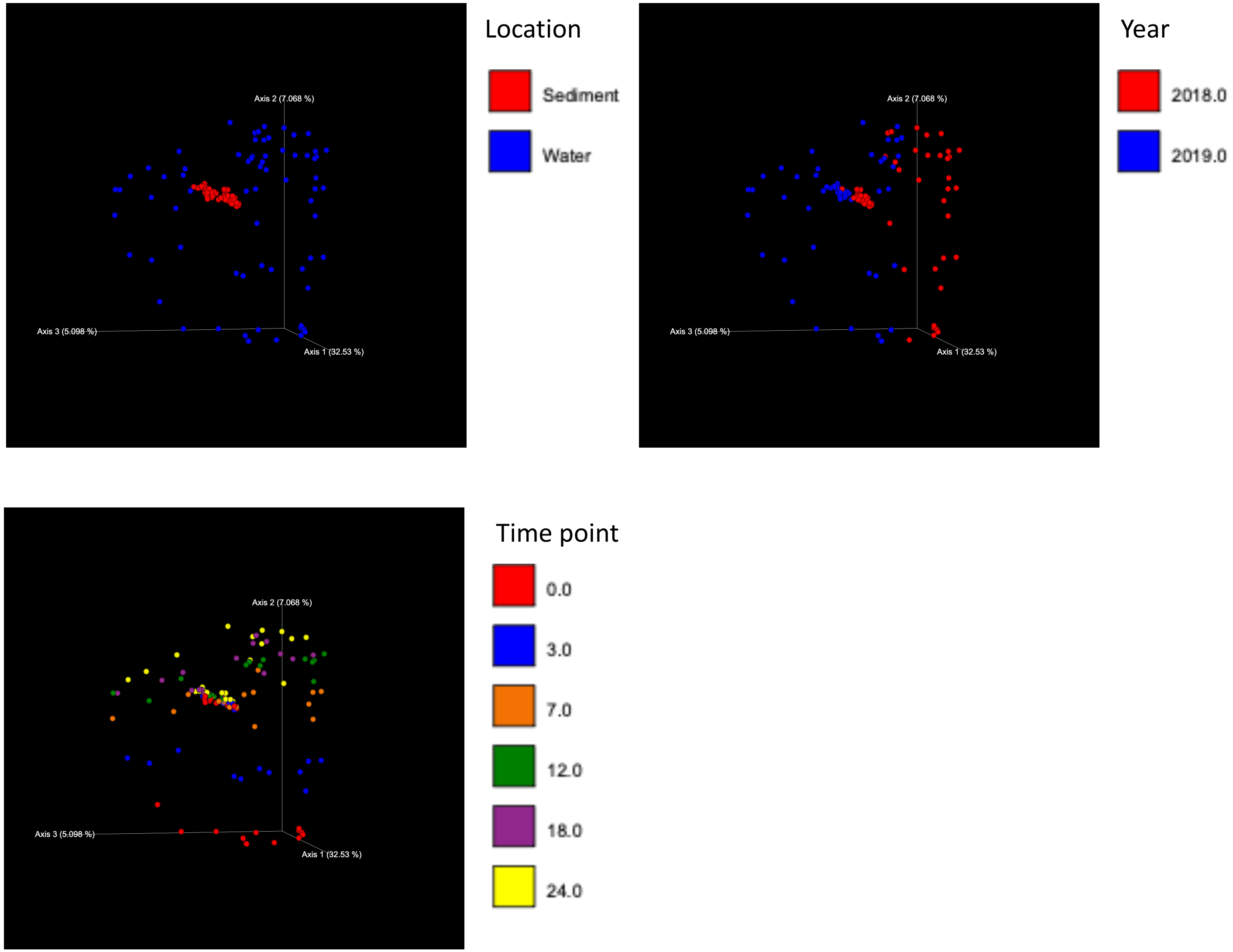


Fig S14. Comparison of 16S rRNA gene diversity by PCoA based on Bray-Curtis dissimilarity between the pilot study (2018) and the main study (2019).
